# From an invariant sensory code to a perceptual categorical code in second somatosensory cortex

**DOI:** 10.1101/2020.03.31.018879

**Authors:** Román Rossi-Pool, Antonio Zainos, Manuel Alvarez, Ranulfo Romo

**Author notes:** Correspondence (R.R.); (R.R.-P.).

## Abstract

A crucial role of cortical networks is the conversion of sensory inputs into perception. In the cortical somatosensory network, neurons of the primary somatosensory cortex (S1) show invariant sensory responses, while frontal lobe neuron responses correlate with the animal’s perceptual behavior. But, where in the cortical somatosensory network are the sensory inputs transformed into perceptual behavior? Here, we report that in the secondary somatosensory cortex (S2), neurons with invariant sensory responses coexist with neurons whose responses correlate with the animal’s perceptual behavior. These distinct neural responses exhibit analogous timescales of intrinsic fluctuations, suggesting that they belong to the same hierarchical processing stage. Furthermore, during a non-demanding control task, the sensory responses remained unaltered while perceptual responses vanished. Conclusively, the S2 population responses exhibit intermediate dynamics between S1 and frontal lobe neurons. These results suggest that the conversion of touch into perception crucially depends on S2.

## INTRODUCTION

Key to understanding the emergence of a percept in the cerebral networks is how sensory inputs are converted into perceptual reports. But, where in the cortical networks do neurons exhibit this conversion? Moreover, is there any cortical area where a sensory representation coexists with a perceptual code? A vibrotactile task in behaving monkeys establishes an appropriate experimental condition to study this problem^1^. While the temporality of each stimulus is represented faithfully and homogenously in the primary somatosensory cortex (S1)^2^, frontal lobe neurons exhibit complex and heterogenous responses associated with working memory and perceptual reports^3^. An intermediate stage between these two extremum processes, in a proposed cortical hierarchy of the cortical somatosensory network, is the secondary somatosensory cortex (S2)^4–6^. Contrary to S1, S2 neurons display large, multi-digit or bimanual receptive fields. Further, their neuronal responses could depend on task context and attention^7,8^, since some show complex activity associated with somatosensory inputs^9,10^. Additionally, previous anatomical evidence has suggested that S2 is largely connected with downstream and upstream areas^11–17^. This means that this single area could have access to faithful sensory inputs, as well as mnemonic information that is solely found in the frontal lobe dynamics. For that reason, we speculate that S2 may play a relevant role in transforming sensory information into perceptual responses. As an extension, we wonder: what is the role of S2 during non-demanding tasks, where perceptual frontal lobe dynamics disappear^1,3,18^? Could this area act as a switch, processing sensory inputs based on task demands? Could the decision-processing observed in frontal lobe areas be ignited by S2 signals? In other words, could S2 act as a gate, permitting transmission of the sensory information based on the task’s requirement?

To address these questions, we analyzed the neural responses recorded in S2, while trained monkeys performed a temporal pattern discrimination task^3^ (TPDT). In each trial, monkeys were asked to decide whether the temporal pattern of two vibrotactile (P1 and P2) stimuli (of equal frequency) are the same or different (Fig. 1A). One of two possible patterns is composed of periodic pulses, and the other is defined by centered grouped pulses, although both patterns last 1 second and maintain the same initial and final pulses. Due to this design, the discrimination is restricted to the period within the boundary pulses. Thus, in order to choose the correct answer (P1=P2 or P1≠P2), the subjects must store the P1 identity in working memory in order to compare it with the identity of P2.

**Figure 1.**
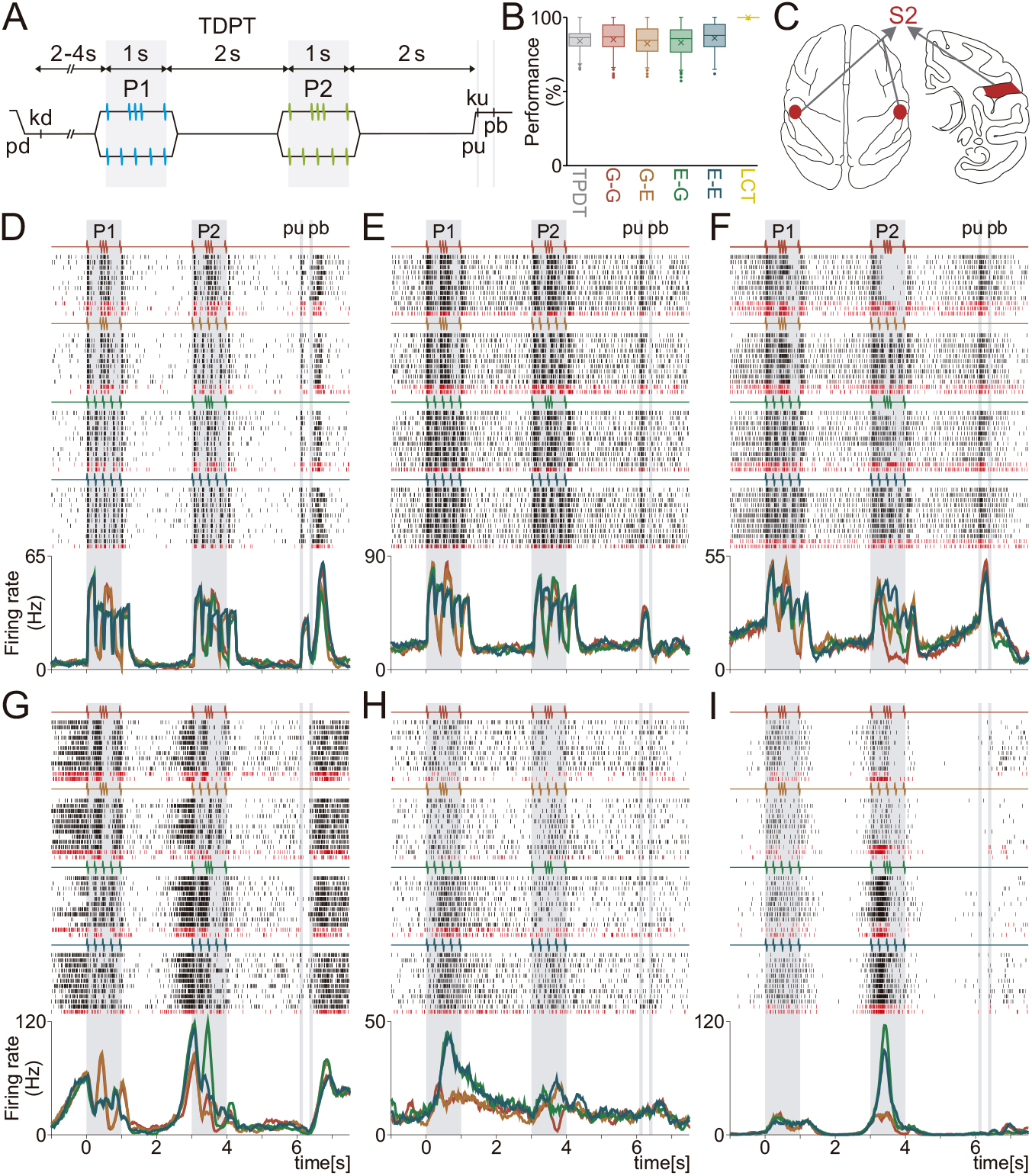
Temporal pattern discrimination task (TPDT) and single neuron activity in S2. (A) Trials’ sequence of events. The mechanical probe is lowered (pd), indenting the glabrous skin of one fingertip of the right, restrained hand (500 μm); in response, the monkey places its free hand on an immovable key (kd). After a variable prestimulus period (from 2 to 4s), the probe vibrates for 1s, generating one of two possible stimulus patterns [P1, either grouped (G) or extended (E); mean frequency of 5 Hz]. Note that in extended pattern (E), pulses are delivered periodically. After a first delay (2s length, from 1 to 3s) between P1 and P2, the second stimulus (P2) is delivered, again either of the two possible patterns [P2, either G or E; 1s duration]; this is also called the comparison period. After a second 2s delay (from 4 to 6s) between the end of P2 and the probe up (pu), the monkey releases the key (ku) and presses, with its free hand, either the lateral or the medial push button (pb) to indicate whether P1 and P2 were the same (P1=P2) or different (P1≠P2). (B) Performance for the whole TPDT (grey, n=423 sessions), for each class [G-G (red), G-E (orange), E-G (green), E-E (blue)] and for the whole LCT (yellow, n=76 sessions) (C) Top (left figurine) of the brain for approaching the second somatosensory cortex (S2) and coronal section of the brain (right) for locations of recordings in S2 (red spots). Recordings were made contralateral and ipsilateral to the stimulated fingertip. (D-I) Raster plots of six S2 neurons sorted according to the four possible classes (stimulus pairs). Each row is a single trial, and each tick is an action potential. Trials were interleaved randomly, although the rows were sorted by class afterward (only 10 out of 20 trials per class are shown). Correct and incorrect trials are indicated by black and dark red ticks, respectively. Average firing rates, per class, demonstrated in traces below raster [peristimulus time histograms (PSTHs)]. Colors distinguish between the four possible classes: G-G (red); G-E (orange); E-G (green); and E-E (blue). Some of the neurons exhibit sensory responses (D-E), others display intermediate (F-G) or categorical activity (H-I).

In previous work we showed that neurons from S1 (specifically, area 3b) faithfully represent the temporal pattern of each stimulus^3^. However, neurons from the dorsal premotor cortex (DPC) coded the different task parameters as broader categories during working memory, comparison and decision periods^3,19,20^. As opposed to S1, the DPC neuronal responses were strongly indicative of the animals’ errors. Further, during a non-demanding task variant (light control task [LCT]), the rich and heterogeneous DPC coding dynamics ceased, while responses from S1 neurons remained unaltered. Based on these results, we hypothesize that an intermediate cortical area between S1 and DPC may act as a switch that depends on the task context, permitting the conversion of the precise sensory inputs into a perceptual categorical response.

Here, we tested this conjecture by analyzing single unit activity in S2, initially identifying exemplary neuronal responses associated with pure sensory, pure categorical, or intermediate dynamics during the cognitively demanding TPDT. Notably, while S2 sensory responses remained unaltered during the LCT, categorical responses ceased. To test this finding within the whole S2 population, we computed the coding dynamics associated with task parameters during the TPDT and LCT. During the TPDT, S2 neurons displayed complex coding associated with the sensory, early working memory, comparison and decision reports. Remarkably, only the sensory coding remains during the LCT. To further investigate these results, we separated the most sensory and perceptual categorical neurons from the whole S2 population. Within the S2 sensory group, we found that the neurons were phase-locked to the stimulus, with dynamics independent of the task context (demanding or non-demanding) and behavioral reports (hits or errors), mirroring the invariant homogeneous responses observed in S1. On the other hand, most of the characteristic responses of the perceptual categorical neurons were severely affected during errors, and entirely ablated during the LCT. While S1 and DPC neurons represent the sensory or perceptual categorical dynamics exclusively, the S2 population reflected a mix of these dynamics, implicating an intermediary role. Furthermore, most of the S2 neurons exhibited responses that mix sensory and categorical dynamics.

To challenge these results, we compared those obtained in S2 with those found in S1 and DPC. The response and coding latencies in the entire S2 population fall between those found for S1 and DPC; however, the S2 sensory neurons reliably exhibited response latencies comparable to those from S1 neurons. On the contrary, the perceptual categorical S2 neurons mirrored the response latencies from the DPC neurons. As an extension, we employed the autocorrelation function in order to estimate the timescales of intrinsic fluctuations in the spiking activity from each cortical area^5^. Notably, population timescales of S2 also fall between those for S1 and DPC, although they were invariant in all cases, even between the subgroups of sensory or perceptual categorical neurons in S2. This means that the neurons commit to completely different dynamics while maintaining a constant inherent timescale. We speculate that this time constant indicates a hierarchical position of the entire S2 network within the perceptual decisionmaking framework; particularly, this position does not depend on the coding dynamics of each subnetwork of neurons. Furthermore, the time constants identified from the autocorrelation functions are not affected during the non-demanding LCT task. Even if S2 and DPC coding dynamics are modified during the LCT, their intrinsic timescales remain unaltered. Collectively, our findings indicate that the S2 network is an intermediate processing area, including neurons with a pure sensory and a perceptual categorical code. Our interpretation is strengthened by the homogenous responses of the S2 sensory neurons, as well as the categorical neurons’ dependence on the monkeys’ behavior and task context (demanding or non-demanding). These results suggest that S2 plays a critical role in the transition from sensory inputs to perceptual behavior.

## RESULTS

### Single neuron responses during the TPDT

We trained two monkeys in the TPDT, in which they reported whether two temporal patterns composed of vibrotactile flutter stimuli (P1 and P2) were the same (P2 = P1) or different (P2 ≠ P1)^3^ (Fig. 1A, Methods). Importantly, the mean frequency (5Hz) was held constant over each pattern’s full duration (1s). The average performance across S2 recording sessions during the TPDT was 84% (±7%), remaining consistent across stimulus pairs or classes (Fig. 1B). We recorded extracellular activity from 1646 neurons in S2 (Fig. 1C) during the monkeys’ performance of the TPDT.

The responses of 15 exemplary S2 neurons are shown in Fig. 1D-I and Fig. S1A-I. Since S2 is typically considered a sensory area, we will denote any sensory-information based response as sensory in order to distinguish from the perceptual categorical responses. Contrary to S1^3^, S2 neurons displayed a broad repertoire of responses, where different types of neuronal dynamics could be recognized. Several neurons were only entrained by the stimulus patterns (Fig. 1D and E; Fig. S1A and B) and their responses faithfully tracked the temporal structure of the stimuli. These neurons were considered sensory and did not exhibit firing rate activity correlated to working memory, comparison or decision reports. Another group of neurons exhibited responses that partially followed the temporal stimulus patterns but were also involved in stimulus transformation and comparison (Fig. 1F and G; Fig. S1C-E). For example, the neuron of Fig. S1E had a much stronger response for grouped patterns (G) during either P1 or P2 stimulus periods. Similarly, the example neuron displayed in Fig. 1F had significantly diminished activity during the comparison for only a specific class (A-A; see Methods where we define class responses); although this neuron distinguished itself in its phase locked response to the stimulus pulses for all other classes (stimulus pairs). During the P2 period, this class selectivity could be associated with each of the four P1 and P2 combinations of stimulus patterns. Therefore, these intermediate neurons exhibit dynamics between pure sensory and pure categorical.

In addition to the intermediate or pure sensory representations, another portion of the S2 population revealed prominently categorical responses. Examples of neurons with these features are shown in Fig. 1H and I and Fig. S1F-I. Notice that these cells do not represent individual stimulus pulses, suggesting that the sensory input is completely transformed into categorical activity, resulting in a more abstract representation related to the identity of P1, P2 or class. Some neurons respond to a specific class selectively with high activity, as illustrated for the example neuron in Fig. S1G. This first panorama of S2 responses was extremely helpful in identifying the most convenient methodological approach to quantify their properties.

### Single neuron responses during the TPDT versus the LCT

Several of the S2 neurons recorded during the TPDT were also measured during the LCT (n = 313), a control variant of the active task. In each trial, the animals received the same stimuli as in the TPDT, but the correct and rewarded reports were guided by a continuous visual cue (Methods). As opposed to the TPDT, the performance for LCT was consistently 100% (Fig. 1B), demonstrating that this task was not comparably cognitively demanding. In a previous work, we observed that neurons in area 3b (S1) do not alter their responses during the LCT^3^. Contrarily, DPC neurons stop coding task parameters during the LCT. Disappearance of DPC coding was observed in single neurons^3^ as well as in their population dynamics^19^. Thus, DPC neurons were recruited to code task parameters (working memory, comparison and categorical perception) exclusively during the TPDT, which means exclusively in a cognitively demanding context.

The examples in Figs. 2 and S2 show the responses of ten typical S2 neurons that were tested in both the TPDT and the LCT. Note that the pure sensory responses are not modified based on task condition (Fig. 2A and S2A), meaning that these neurons always communicate sensory information; the invariance demonstrated in these neurons directly parallels the responses that have been observed in S1 (area 3b). On the other hand, neurons that depict a mix between sensory and categorical responses do alter their coding dynamics somewhat during the LCT (Fig. 2B and D, Fig. S2B, C, E and F). For example, the neuron shown in Fig. S2C displays a strong affect in its response. Whereas this neuron exhibits G-pattern differential responses during P2 in the TPDT, the neuron has no categorical response during the LCT. Neurons with strong categorical responses (Fig. 2C and S2D) stop coding task parameters by diminishing their response during the LCT. Clearly, the activity from these neurons is drastically modified during the LCT. While some S2 neurons change their dynamics completely during the LCT, the response of other subgroups of neurons remain partially or completely unaltered in both task contexts.

**Figure 2.**
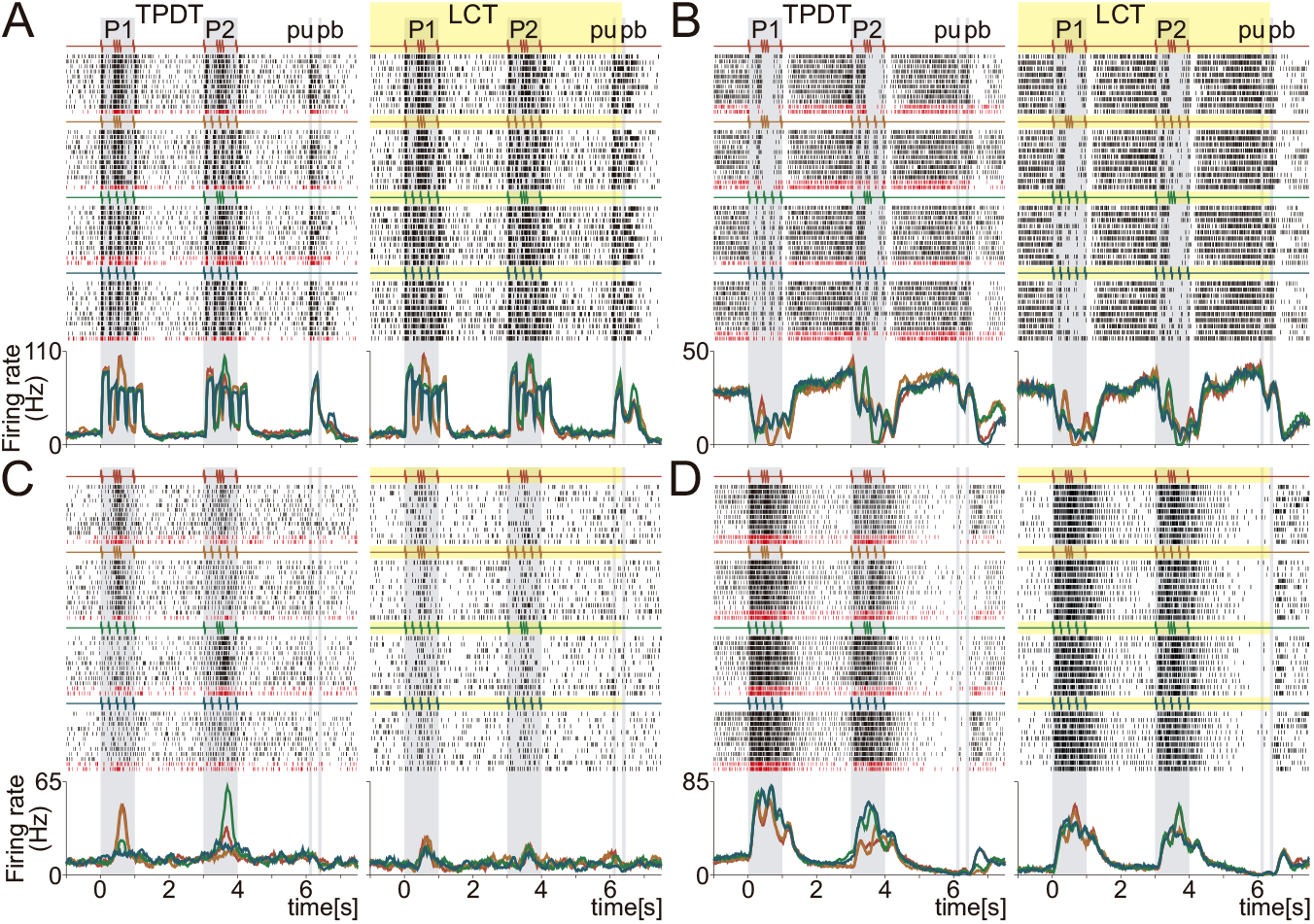
Single S2 neurons activity during the TPDT and LCT. (A-D) Raster plots of four S2 neurons tested in both tasks: TPDT (left) and LCT (right). Response rows sorted by correct and incorrect trials, row blocks sorted according to the four possible classes delivered during P1 and P2 (stimulus pattern pairs). Correct and incorrect trials (black and red ticks, respectively) only for the TPDT, since there were no errors during LCT. Traces below the raster plots are class average firing rates per neuron and condition. Each color refers to one of the four classes: G-G (red); G-E (orange); E-G (green); and E-E (blue). These neurons are different from those in Fig. 1. Sensory responses invariant during TPDT and LCT (A, B and D), while categorical responses ceased during the LCT (C).

### Context-dependent coding dynamics

To measure the coding capacities of S2 neurons as a function of time, we employed ROC (receiving operating characteristic) comparisons between the two firing rate distributions, where each was conditioned to a single stimulus class: G-G, G-E, E-G and E-E. Then, each time bin was tested for identification of one of the four coding profiles associated with different task parameters: stimulus identity for each presented pattern (P1 or P2), class selectivity for each pair of patterns, or decision outcome (Methods). Through the application of this procedure, we were able to calculate the percentage of S2 neurons (n=1646) that coded each task parameter during the TPDT (Fig. 3A). With respect to P1, a large percentage of S2 neurons coded the identity of the pattern (cyan colour) during the P1 period. This means that the identity of P1 could be decoded from an abundant number of cells while employing a 200 ms firing rate window. Remarkably, the number of S2 neurons coding P1 identity decreases significantly at the beginning of the working memory period of the TPDT (between the end of P1 and the beginning of P2). The populational decrease in P1 representation is not absolute (~5% remain), but it does demonstrate that the P1 identity correlate is mostly limited to the P1 stimulation period and the initial component of the working memory period. Additionally, when each single neuron’s dynamics was analysed individually, no S2 neuron demonstrated persistent P1 coding. The remaining 5% is a result of the overlap of the early and late P1 coding dynamics. This result corroborates previous works suggesting that persistent coding is negligible in S2 neurons^2,21,22^. Remarkably, at the end of the working memory period and the beginning of P2, the percentage of S2 neurons with P1 coding increased. This reappearance in P1 coding is mainly due to late neurons, as the one shown in Fig. 1I. Even though P1 occurred 2s earlier and the information of P1 is negligibly maintained throughout the delay period, the response of several S2 neurons became strongly modulated by P1. A possible role for these neurons is reinvocation of P1 information for use within the network during the comparison period (P2 = P1 or P2 ≠ P1), also known as the P2 stimulation period.

**Figure 3.**
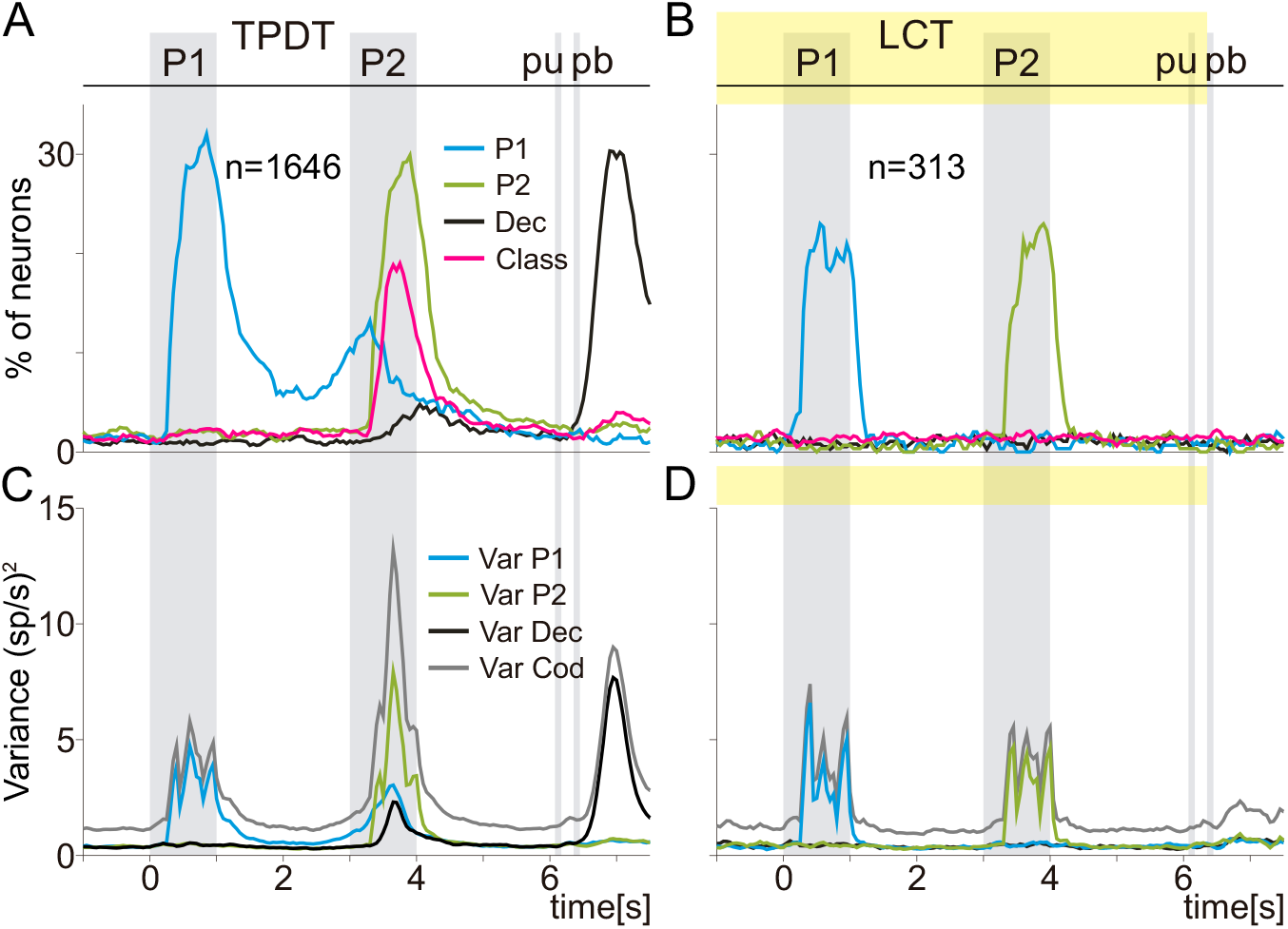
Population coding dynamics and population variance in S2 during TPDT and LCT. (A-B) Percentage of neurons with significant coding as a function of time during TPDT (n=1646) and LCT (n=313). Traces refer to P1 (cyan), P2 (green), all class coding (pink, stimulus pattern pairs), and decision coding (black). (C-D) Population instantaneous coding variance (*Var_COD_*, grey, Eq. 1), P1 variance (*Var_P1_*, cyan, Eq. 2), P2 variance (*Var_P2_*, green, Eq. 3), and decision variance (*Var_Dec_*, black, Eq. 4) during TPDT (right, n=1646) or LCT (left, n=313). Note that P1 working memory, decision, and class coding, as well as variances, essentially vanished during LCT: coding restricted to stimulus periods in LCT. For details, see Methods.

The comparison periods (from 3 to 4s) began with the extinction of the P1 working memory signal, in concordance with the appearance of a high percentage of neurons coding P2 (green trace). The number of neurons coding P2 is akin to that observed during P1. On the other hand, a strikingly large number of neurons with class selective coding (pink trace) emerged almost simultaneously with P2 coding. Class coding applies to neurons with a categorical response associated to any of the four possible P1 and P2 combinations. For instance, neurons depicted in Fig. 1F and S1G are both selective to class G-G. The S2 neurons with a class coding signal began during P2 and decreased thereafter, reaching extinction slightly before the P2 signal. Besides coding for P1 stimulus identity and classes during the P2 period (comparison period), a small percentage of S2 neurons also exhibited the decision outcome (either P2=P1 or P2≠P1). Surprisingly, the small percentage evolves into a massive portion of S2 neurons reflecting the monkeys’ categorical decision during the motor report period. Although the particular role of this decision signal is unclear, it was also observed in DPC during the same period and task^3^.

To what extent are those S2 signals dependent on the animal’s judgment? Are these coding dynamics modified during the non-cognitively demanding task (LCT)? To answer these questions, we applied the same decoding scheme [based on receiving characteristic analysis (ROC analysis)] to the population of S2 neurons recorded during the LCT (n=313). Remarkably, coding dynamics changed dramatically during this task (Fig. 3B). Under this change in context, S2 neurons coded only information related to the actual stimulus pattern. Hence, the S2 dynamics during LCT are exclusively sensory, exemplified by neurons in Fig. 2A and B. In further comparison to the TPDT, the total percentage of neurons coding P1 and P2 identity each decreased during the LCT. Notice that early and late coding during the working memory period is abolished. Similarly, class coding vanished fully during the comparison period (see neurons in Fig. 2C and Fig. S2D). Furthermore, note that the strong categorical decision signal, observed in TPDT after pu, also disappeared (see neuron in Fig. S2E and F). As a result, S2 activity has a profound relationship with the behavioral context, as well as the cognitive processing of the stimuli.

To further quantify said relationship, we computed the response’s variability associated with the task parameter coding, employing the population instantaneous coding variance (*Var_COD_*, Eq. 1, grey traces) during the TPDT (Fig. 3C) and the LCT (Fig. 3D). This measure estimated, at each time bin, the activity fluctuations between classes and neurons. When more neurons code task parameters, higher values of *Var_COD_* are generated^20^. To elaborate upon this analysis, we calculated the population variances related to P1 (*Var_P1_*, Eq. 2, cyan traces), P2 (*Var_P2_*, Eq. 3, green traces) and categorical decision outcome (*Var_Dec_*, Eq. 4, black traces) in both task conditions. There was high coincidence between the effects observed with these different measures and the corresponding coding dynamics (Fig. 3A-B). While coding variances wax and wane during the working memory period in the TPDT, they ceased in the LCT. The most drastic change occurred at the comparison period: during this period in the TPDT, *Var_COD_* reached its maximum value while single neurons exhibited a high diversity in coding responses (Fig. 3C). Opposingly, the coding variance was only related to the P1 and P2 stimulus identity during the LCT. Note the large difference between the maximum value of *Var_COD_* during the TPDT (~13(sp/s)^2^) and the LCT (~5(sp/s)^2^). During the TPDT, the variance related to decision outcome (*Var_Dec_*) exhibits high values after pu, but this dynamic vanished during the LCT. These findings substantiate our claims regarding S2 responses; the responses associated to perceptual task parameters are strongly dependent on the task’s context.

### Phase-lock and categorical neurons

Based on the above findings, we aimed to recognize S2 sensory neurons whose evoked spikes were phase-locked to the stimulus pulses. When a small window is employed to compute the firing rate (24 ms, Methods), the response of a sensory neuron (Fig. 4A) during the stimulus periods should be periodic during E-pattern (golden trace) but aperiodic during G-pattern (green trace). This difference in periodicity is displayed by the example of Fig. 4A, which illustrates the typical phase-locked response during P1. To estimate the degree of periodicity of individual neurons, we computed the power spectrum this periodic E-pattern frequency, during the stimulation of each trial (Methods). Actually, the inherent E-pattern periodic frequency is 4.16Hz. In trials with periodic pattern, this power spectrum value depends on the periodicity of the spike trains. As such, in neurons with phase-locked responses, the power spectrum of this frequency should give a high amount of information about the stimulus identity^2^. In other words, it should be possible to decode pattern identity based on the spike train periodicity in sensory neurons. We calculated this periodicity information (*I_Per_*, Eq. 10) for each neuron during P1 and P2. Analogous results were found using both stimulation periods, so we chose to show P1 results in the rest of this work. Whereas the periodicity information is high in the example of Fig. 4A (*I_Per_*=0.89Bits), this metric is low in categorical neurons without phase-locked responses (Fig. 4B, *I_Per_*=0.04Bits). This means that this information value allows us to recognize neurons with strong phase-locking to the temporal stimulus pattern. Based on that, neurons with significant and high values of *I_Per_* (>0.25Bits) were classified as sensory. This value was set to isolate the most sensory neurons from the whole population to study their dynamics in detail. Even if it is an arbitrary separation, it serves to detect specific response features of these neurons.

**Figure 4.**
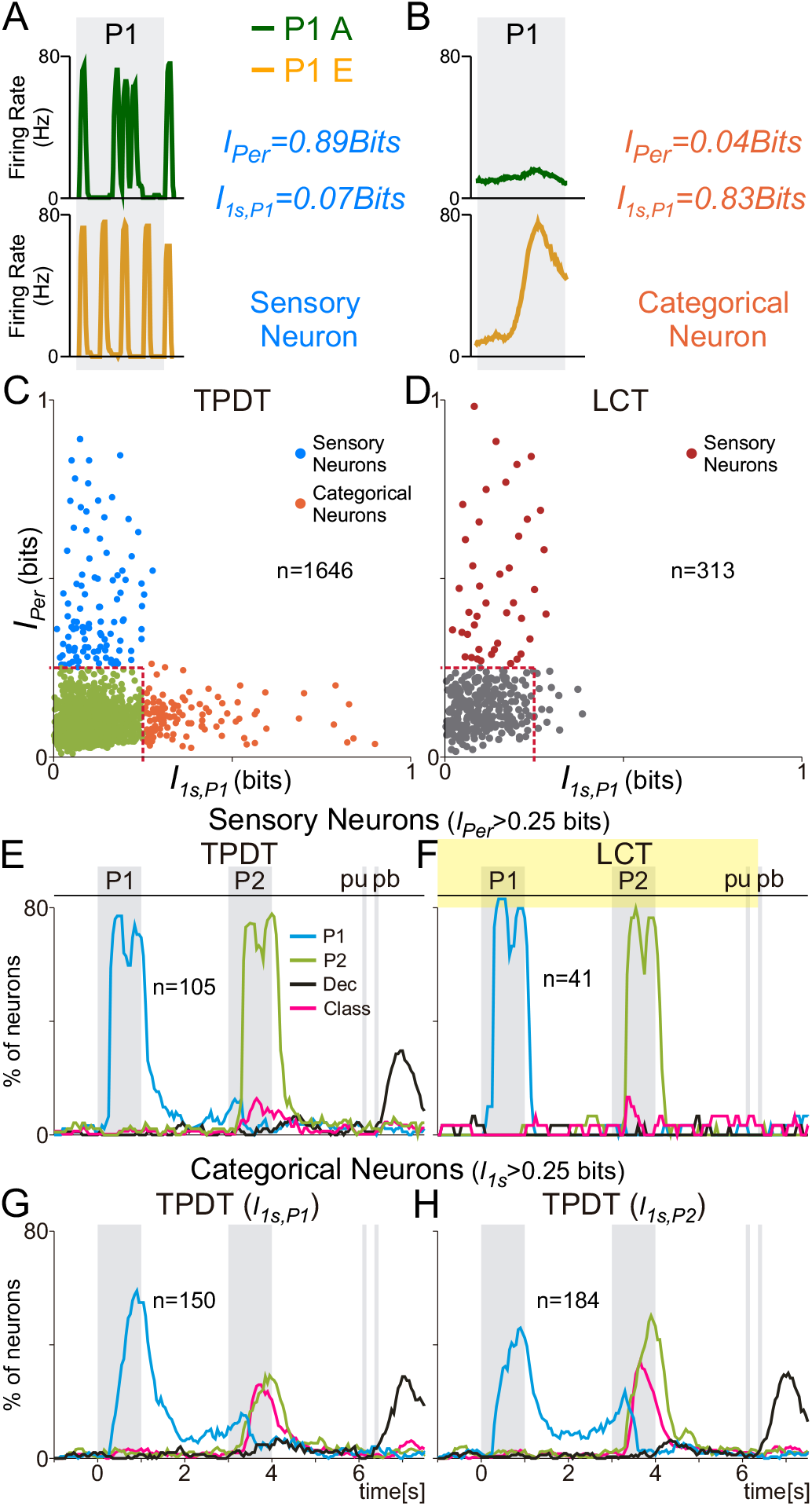
Sensory and categorical neurons and their coding dynamics. Periodicity mutual information (*I_Per_*, Eq. 10) and 1s firing rate mutual information (*I_1s,P1_*, Eq. 6) associated with pattern identity; used to identify sensory and categorical S2 neuron subpopulations. (A) Exemplary S2 neuron with high periodicity information during the first stimulation period (*I_Per_* 0.89bits). This neuron depicts low 1s firing rate mutual information associated with P1 identity (*I_1s,P1_*=0.07bits). (B) S2 neuron with low periodicity (*I_1s,P1_*.0.04bits) and marked categorical response for E pattern. This neuron conveys large 1s firing rate mutual information related to P1 identity (*I_1s,P1_*=0.83bits). (C) For each S2 neuron recorded during TPDT (n=1646), 1s firing rate mutual information (*I_1s,P1_*, x axis) is plotted against periodicity mutual information (*I_Per_*, y axis), both associated with the identity of the first stimulus pattern (P1). P2 results analogous so not depicted. The red dashed lines indicated the mutual information limit (*I*>0.25bits) used to label S2 neurons as sensory or categorical. These are arbitrary boundaries, although useful to explore the dynamics features of the S2 neurons with higher amounts of *I_Per_* or *I_1s,P1_*. Most S2 neurons exhibit low or intermediate values for both *I_1s,P1_* and *I_Per_* (green points). (D) *I_1s,P1_* (x axis) is plotted against *I_Per_* (y axis) for each S2 neuron recorded during the light control task (n=313). Note that much fewer neurons exhibit *I_1s,P1_*>0.25bits during LCT. (E-H) Percentage of each subpopulation of neurons with significant coding as function of time during TPDT or LCT. Traces refer to P1 (cyan), P2 (green), class (pink, stimulus pair combinations), and decision coding (black). (E-F) Sensory neurons (*I_Per_*>0.25bits) during TPDT (right, n=105) or LCT (left, n=41). Most sensory neurons only involved in coding P1 or P2 identity during stimulation periods. (G-H) Categorical neurons computed during the first stimulus (*I_1s,P1_*>0.25bits, n=150) or second stimulus (*I_1s,P1_*>0.25bits, n=184) periods. In both cases, P1 coding emerges later than in sensory neurons, although maintained for a longer period; class coding is present in both types of categorical neurons. Decision coding after “pu” is observed in both categorical and sensory neurons during TPDT.

To separate neurons with categorical responses instead, we computed the 1s firing rate mutual information associated with the identity of P1 (*I_1s,P1_*, Eq. 6) or P2 (*I_1s,P2_*). These measures used a 1s window to calculate the firing rate of each trial during the whole stimulation period. Consequently, the temporal structure of the spikes evoked during stimulus periods during each trial is irrelevant to this metric; it condenses all responses into a single firing rate calculation. By this we mean that phase-locked responses to the stimuli will have approximately the same number of spikes during E or G patterns (Fig. 4A). Contrary to that, neurons with strong categorical and differential responses will have different values of activity for each specific pattern (Fig. 4B). Hence, high values of *I_1s,P1_* mean that the neuron responds differentially for a specific pattern, in this case P1. The differential response to E-pattern shown in Fig. 4B gives rise to a high value of *I_1s,P1_* (0.83Bits). Intuitively, the sensory neurons (Fig. 4A) produce minimal amounts of *I_1s,P1_* (0.07Bits). Neurons with significant and high values of *I_1s,P1_* (>0.25Bits) were labeled as categorical responses. Again, although the information value used to classify sensory and categorical responses (*I* >0.25Bits) was arbitrary, the intent was to detect neurons with extreme responses in the whole population. The resultant subpopulations yield valuable and distinct results, demonstrated analytically in the following sections.

Each point in Fig. 4C represents a single neuron recorded during the TPDT (n=1646), with the position in the plot defined by *I_1s,P1_* (x axis) and *I_Per_* (y axis). Notice that a comparable number of sensory (n=105) and categorical (n=150) neurons were identified with our criteria (*I* >0.25Bits). Besides, the neurons not classified as sensory or categorial represented the brunt of the population, exhibiting low or intermediate values for both types of information (green points). Thus, this criterion was effective in selecting a comparable number of neurons with the purest categorical or sensory responses. Similar subgroups of neurons were found with a much more stringent criterion, *p<0.003* instead of *p<0.01*. Further, similar cumulative distributions for *I_Per_, I_1s,P1_* and *I_1s,P2_* (Fig S3A) were found during the TPDT. Thus, the fraction of neurons with information smaller or equal to the amount indicated on the *x* axis is comparable between metrics. Remarkably, when using the same metrics for the neurons recorded during the LCT (n=313, Fig. 4D and Fig S3B), the plots exhibited large changes. In the LCT, almost all high values of mutual information were with respect to periodicity, so they were identified as sensory neurons (Fig. 4D). Therefore, the categorical neurons decreased drastically during the LCT. The same trend was found with the cumulative distribution (Fig S3B): higher values of *I_Per_* are much more common than *I_1s,P1_* and *I_1s,P2_*. These results support those observed in Fig. 3 with the variance and the coding dynamics.

### Sensory versus categorical coding dynamics

To analyze the properties of the dynamics for each subgroup of neurons, we implemented the same coding scheme as in Fig. 3. The coding dynamic observed in the sensory population during the TPDT (Fig. 4E) changed appreciably with respect to the whole population (Fig. 3A). In contrast to Fig. 3A, during both stimulus periods, the coding dynamics of sensory neurons increased abruptly; P1 and P2 coding evolved similarly during both stimulus periods. Notice that the percentages with significant coding during P1 or P2 periods were much higher in Fig. 4E than Fig. 3A (80% versus 30%). Further, when we computed the same coding for to the LCT sensory neurons (n=41), an akin response was found (Fig. 4F); both subpopulations attain similar percentages of pattern identity coding neurons, with analogous dynamics. These findings contrast sharply with the differences observed between the TPDT and the LCT (Fig. 3) for the whole population. Note that the categorical decision signals after pu were still present in several sensory neurons during the TPDT, but not during the LCT. Likewise, applying variance measures to these subpopulations of sensory neurons produced similar results (Fig. S3C-D). Except for the categorical decision after pu, variance responses are analogous between TPDT and LCT. Thus, variance dynamics are restricted to stimulation periods in sensory neurons during both task conditions. Notice that the values of variance observed in these highly informative neurons (Fig. S3C-D, ~30(sp/s)^2^) were much larger than in the whole population (Fig. 3C-D, ~10(sp/s)^2^).

Neurons that convey high values of categorical information associated with P1 or P2 identity (*I_1s,P1_*>0.25Bits, n=150 or *I_1s,P2_*>0.25Bits, n=184) gave rise to different coding dynamics (Fig. 4G-H) in comparison to sensory neurons (Fig. 4E). Again, the percentage of neurons with P1 or P2 identity coding is higher than in the whole population (Fig. 3A). Note that the increment of P1 or P2 coding during stimulus periods is not as abrupt as in Fig. 4E. Additionally, several neurons code P1 identity during the early part of the working memory period. In expected contrast to the sensory coding dynamics (Fig. 3A), categorical neurons display a high percentage of class coding neurons (pink trace, Fig. 3G-H). The variance in the responses of these categorical subgroups (Fig. S3E-F) yielded similar features. Contrastingly, these measures increased much more slowly during P1 or P2 than in sensory neurons. Particularly, categorical neurons identified during P2 (Fig. S3F) showed elevated values during the P2 period. This means that these neurons take a preponderant role during the comparison period of the TPDT.

### Single neuron periodicity and categorical information between task contexts

To investigate whether mutual information values depended on the cognitive context (TPDT or LCT), we compared the same metrics (*I_Per_* and *I_1s,P1_*) in a subgroup of neurons (n=313) recorded during both TPDT and LCT. Based on the differences observed between the TPDT and the LCT responses (Fig. 3), we speculated that single neurons may change their information values between these task conditions. Each point in Fig. S4A was established by two values, calculated on the same neuron: the TPDT *I_Per_* (x axis, Eq. 10) and the LCT *I_Per_* (y axis). Remarkably, S2 neurons exhibited a trend to have higher periodicity information values during the LCT. The angle distribution was biased to higher values (<θ>=57.49°). Thus, neurons with a higher degree of phase-locking are more common during the LCT than the TPDT.

In contrast, neurons displayed larger values of categorical information (*I_1s,P1_*) during the TPDT than in the LCT (Fig. S4B). In other words, during the cognitively demanding task (TPDT), the S2 neuron responses conveyed higher information associated with pattern identity when we integrated their activity during the whole stimulation period. In concordance, the angle distribution was biased to low values (<θ>=31.45°). Thus, whereas the periodicity information tends to increase during the non-demanding task (LCT), the categorical information has the opposite effect. Single neurons exhibit a tendency to be more sensory during the control task, but more abstract and categorical during the demanding task. These findings are in concordance with Fig. 3, where sensory responses were more common during the LCT, whereas categorical coding was more abundant during the TPDT. Consequently, this strengthens the argument regarding S2 playing a switching role in the cortical network.

### Sensory population responses during hits and error trials

In this section, we focus again on the subpopulation of sensory neurons (n=105). For each of the four classes of stimulus pairs, the firing patterns of the example neurons (Fig. 1D-F) were like those observed in the normalized population activity during hit trials (Fig. 5A, Methods). Notably, population average responses were entrained to the vibrotactile stimulus patterns, but beyond that, there was no sign of any firing rate modulation associated with the working memory or comparison periods of the TPDT. Therefore, in general, the sensory S2 spike trains faithfully tracked the internal temporal structure of the stimulus patterns. To further document their sensory responses, we compared the response patterns evoked during hits (Fig. 5A) and error trials (Fig. 5B). We found no statistical differences between the respective responses based on mean squared errors (variations of 1.2–2.3 % for each stimulus class). These findings indicated that there were no significant differences among sensory responses between hits and errors for any stimulus class. Moreover, the average sensory response was indistinguishable from the average sensory response in the LCT (Fig. 5C, n=41). Thus, we conclude that S2 sensory neurons encode the stimulus patterns accurately, but only during the stimulation periods, regardless of task context and the monkey’s performance.

**Figure 5.**
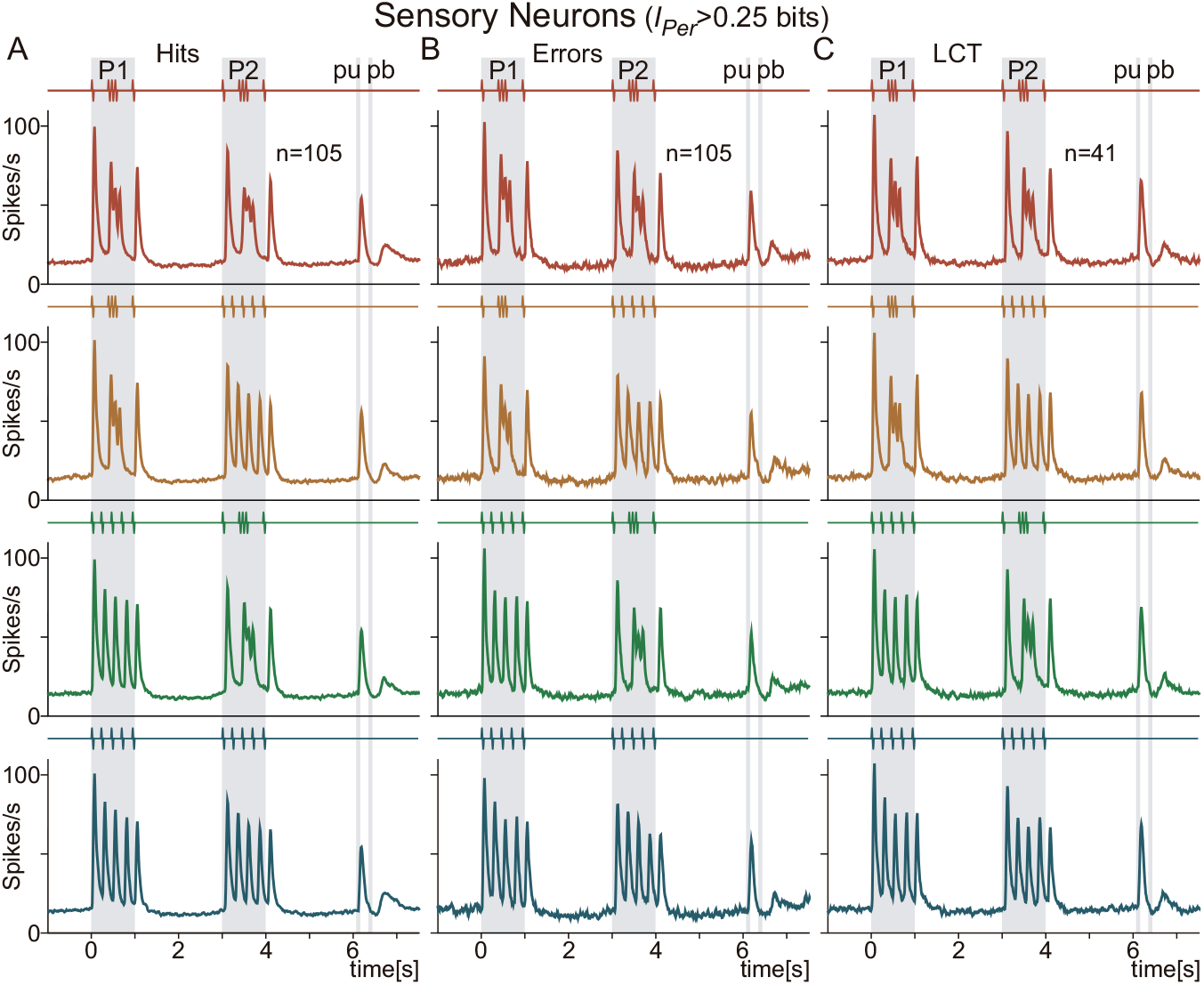
Sensory responses during hits, errors and LCT trials. Sensory S2 neurons selected when periodicity information is significant and higher than 0.25 bits (*I_Per_*>0.25 bits, Eq. 10). (A) Normalized S2 sensory population activity for hits, (B) errors (n=105), and (C) LCT (n=41). Even if the number of neurons is the same for hit and error normalized sensory population responses (n=105), the number of error trials is far fewer. For each trial class, differences between classes or conditions calculated using integral square error; found to be quite small (from 1.2 to 2.3 %).

### Population P1 coding during hit and error trials

To investigate the degree to which the activity in S2 neuron responses correlated with the monkey’s choice, we examined whether the firing rate mutual information associated with P1 identity was somehow different between correct (hit) versus incorrect (error) trials. For this, we first used the normalized activity (z-score) at each time bin from the 1253 neurons with significant P1 coding (Methods). Note that in this case we computed the activity from each neuron using a 200 ms sliding window displaced every 50 ms. Then, we split the normalized responses into hit and error trials and measured the P1 mutual information employing the z-scored responses across stimulus identity within each group (*I_P1_(t)*, Eq. 7). Note that while *I_1s,P1_* (Eq. 6) computed the information at a single time bin that covers the whole stimulus period, *I_P1_*(*t*) (Eq. 7) measured the information associated with P1 as a function of time, employing 200 ms windows. For the neurons with P1 coding (n=1253), we previously determined that the responses associated with G and E patterns were very different, and thus highly informative, during hit trials across time. This result was further confirmed with *I_P1_*(*t*) for hit trials (Fig. 6A, blue trace). In this figure, we concentrated on the first part of the task, from 0 to 3s. Note that S2 neurons display elevated values of *I_P1_*(*t*) during the early part of working memory. Even if neurons still convey P1 information during error trials (Fig. 6A, red trace), its value decreased significantly during the last part of the stimulus and the working memory period. Thus, when the monkey made an error, S2 neurons demonstrated diminished information carried about P1 identity.

**Figure 6.**
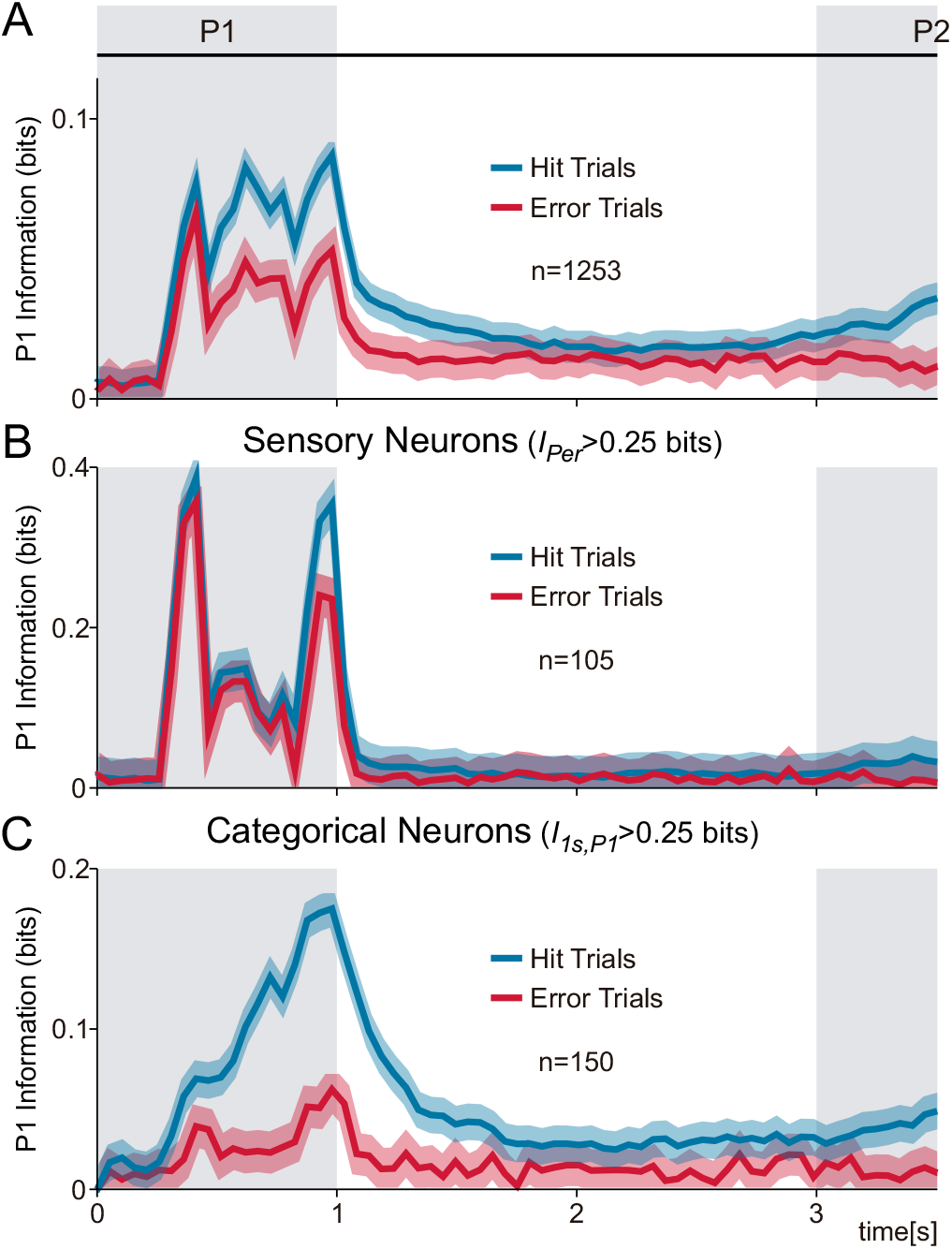
P1 Mutual information in sensory versus categorical neurons during hit and error trials. S2 population firing rate mutual information associated with P1 identity, computed as a function of time (*I_P1_(t)*, Eq. 7) during hit (blue) and error (red) TPDT trials. (A) Neurons with at least 4 consecutive time bins with significant P1 coding (n=1253) were employed to calculate *I_P1_(t)* during hit and error trials. (B) Sensory S2 neurons (*I_Per_*>0.25 bits, Eq. 10, n=105) employed to estimate *I_P1_(t)*. Notably, mutual information nearly unaltered during error trials. (C) Categorical S2 neurons (*I_1sP1_* >0.25bits, Eq. 6, n=150) were used to calculate *I_P1_(t)* in hits and errors. Notably, hits mutual information increased slower than in sensory neurons, and *I_P1_(t)* remained small during error trials. Shadows indicate the information confidence intervals at 95%.

The previous graph obscures the nature of the S2 responses, whereby neurons convey P1 information with different response features. In Fig. 6B, we show the P1 mutual information as a function of time (*I_P1_*(*t*)) for the sensory subpopulation of neurons (*I_Per_* >0.25Bits, n=105), employed previously to compute Figs. 4E and 5. In agreement with Fig. 5, sensory neurons conveyed information analogously during hit (blue trace) and error trials (red trace). Further, these neurons only coded sensory inputs during the stimulation period.

Nonetheless, in the case of categorical neurons (*I_1s,P1_*>0.25Bits, n=150), the results contrast drastically with the previous figures. Markedly, *I_P1_*(*t*) increases in a slower manner during hit trials in this subpopulation of neurons (Fig. 6C, blue trace). This means that categorical information emerged later than sensory, and the most informative point was at the end of the stimulation period. Further, contrary to sensory neurons, categorical neurons conveyed P1 information during the early part of the working memory period. Most notably, the pattern identity information carried by categorical neurons almost vanished during error trials (Fig. 6C, red trace). This means that the P1 identity is no longer coded during error trials. Hence, this subgroup of S2 neurons are more related to the behavioural responses. In conclusion, while activity from sensory neurons do not covary with monkeys’ behaviour, categorical responses are strongly linked to it.

### Categorical decision coding during hit and error trials

To evaluate how the categorical decision signal unfolded over time, we analyzed the neurons with significant decision coding (n=721) during hit and error trials. We computed a population choice probability index (CPI, Methods, Fig. S5A) over time to compare the distributions for hit versus error trials in classes associated with identical decision outcome (c1 and c4 for P2 = P1; and c2 and c3 for P2 ≠ P1). Note that in this case we focus on the last part of the task (from 3s to 7.7s), where we found similar CPI evolution for match (P2=P1, green trace) or non-match trials (P2≠P1, red trace). Even if CPIs reach significant values at the end of the comparison periods (CPI~0.6), higher values were observed after the push button press (CPI~0.8). This strong decision signal agreed with the dynamics observed in Fig. 3. Similar as we did in Fig. 6, we also computed the 200 ms-firing rate mutual information associated with the categorical decision during hit and error trials (Eq. 8, *I_Dec_(t)*, Fig. S5B). The results were consistent with those observed in Fig. S5A: it is possible to decode the categorical decision identity from either hit (blue trace) or error trials (red trace).

Next, we estimated the 200 ms-firing rate mutual information associated with reward (Eq. 9, *I_Rew_(t)*, Fig. S5C). To compute this, we divided the z-score responses into two distributions: hit and error trials. This means that we have one distribution that combined the whole population responses during hit trials and another distribution for error trials. Note that these distributions are different from the used in Fig. S5B, where trials were first split according to the stimulus pair. Notably, S2 neurons carried significant reward information during the period after pb press (after 7.5s, Fig. S5C). Hence, it is possible to employ S2 activity to infer if the animal received reward or not. Note that this signal emerged subsequently to the categorical decision signal that appeared after pb. Even if the relevance of this coding is not evident, we speculate that it could be crucial to develop the S2 dynamics observed during the TPDT.

### Response and coding latencies

We hypothesized that the different subpopulations of S2 neurons will respond to and code P1 with different response latencies. For each neuron from the whole population of S2 (n=1646) recorded during TPDT, we first calculated the response latency: the initial time at which its activity increased significantly above baseline (Fig. 7A, green, Methods). In Fig. 7A, we show the response latency probability distribution for the whole S2 population, displaying a mean value of 97 ± 94 ms (± standard deviation, S.D.). These measures computed how quickly the S2 neurons began to respond, but not when they began to code the stimulus pattern. Then, we calculated the coding latency probability distribution for S2 as whole (Fig. 7A, orange, Methods). Expectedly, the coding latencies (mean: 432 ± 232ms) exhibited much longer values than those for the response latencies (AUROC *p<0.001*). Further, S2 sensory neurons (*I_Per_*>0.25Bits, n=105, Fig. 7B) were significantly (*p<0.001*) shorter in response (33 ± 15ms, pink) and coding latencies (301±71ms, blue). Thus, the sensory subgroup is evidently the fastest portion of S2 neurons, both in response to and codification of the stimulus identity. The opposite tendency was observed among categorical neurons (*I_1s,P1_*>0.25Bits, n=150, Fig. 7C). In comparison, this subgroup of neurons showed much longer response (106 ± 15ms, dark red) and coding (477 ± 201ms, light blue) latencies than the whole S2 population (*p<0.005*). Markedly, there is a significant difference between the categorical and the sensory latency distributions (compare Fig. 7B with 7C; *p<0.001*). Further, the whole population recorded during the LCT (n=313, Fig. 7D) display a trend to be faster than the TPDT population (*p<0.05*): mean value for the response latency was (72 ± 88ms) and for coding latency was (363 ± 170ms). To further test these differences, we computed for each population the cumulative probability for the response latency (Fig. 7E) and for the coding latency (Fig. 7F). Again, sensory neurons were the fastest and categorical the slowest for both types of latency. Even if the differences between the TPDT and the LCT were not large, a clearer tendency could be observed in the cumulative curves (Fig. 7F). This result is in concordance with Fig. S4 that shows that S2 neurons are more sensory and less categorical during the LCT.

**Figure 7.**
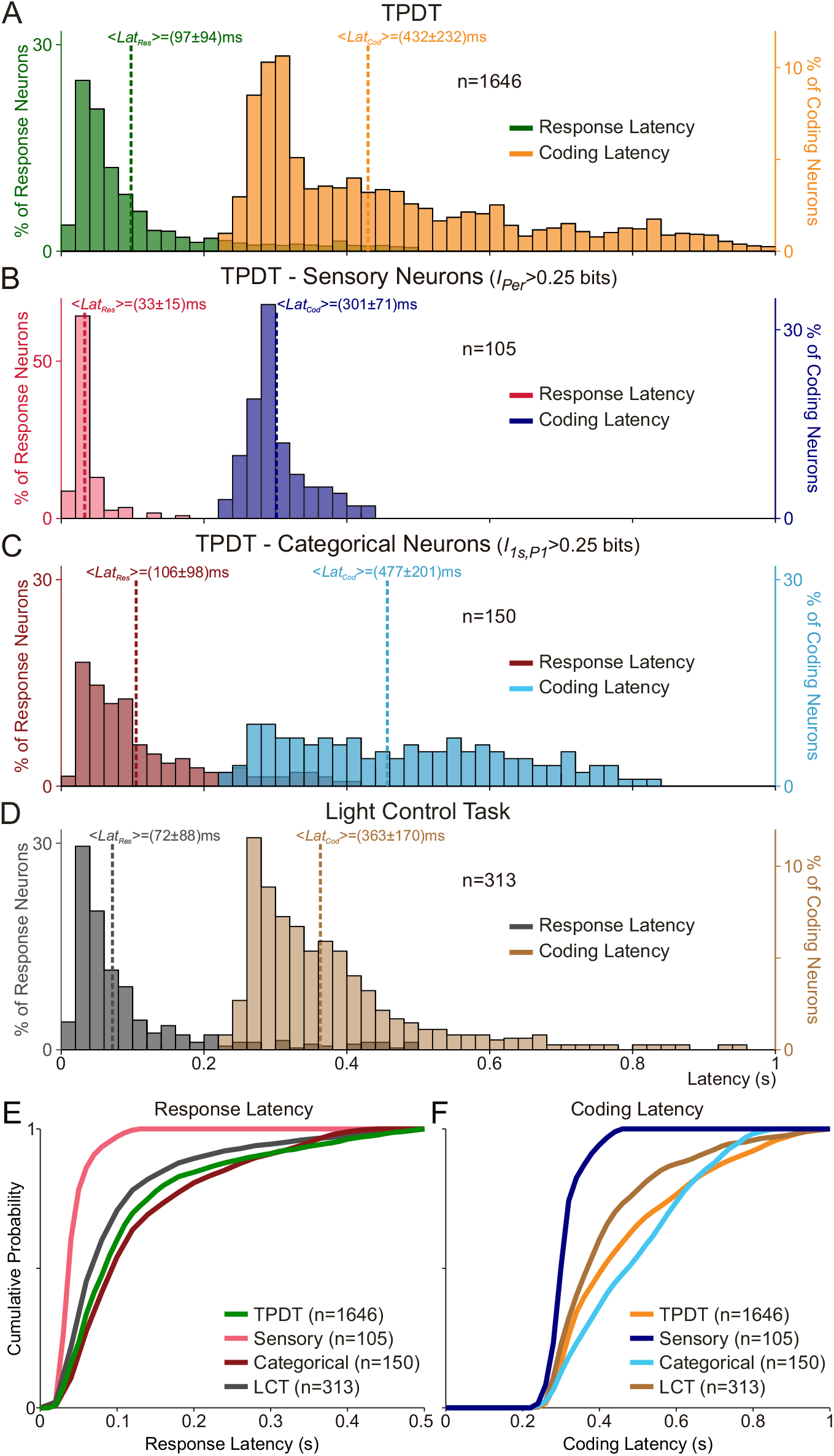
Response and coding latencies in the S2 population. (A-D) Response (*Lat_Res_*, left axis) and coding latency (*Lat_Cod_*, right axis) distributions are computed for: (A) the whole population of TPDT neurons (n=1646); (B) sensory neurons during TPDT (n=105); (C) categorical neurons during TPDT (n=150); (D) the whole population of LCT neurons (n=313). Mean values for each latency distribution are indicated with dashed lines (± values indicate standard deviation, S.D.). (E-F) Cumulative probability distribution for response and coding latency. The value on the y axis represents the fraction of neurons with response and coding latency smaller or equal to the value indicated on the x axis.

To further explore the role of the S2 population in the somatosensory hierarchy, we recruited previously analyzed cortex populations (area 3b, S1 and dorsal premotor cortex [DPC]) for comparison between both of their latency probability distributions (Fig. S6). In both cases, 3b neurons were fastest (blue color, auROC *p<0.001*). Electrophysiological evidence suggests that 3b neurons constitute the earliest somatosensory information input to the cortex^2,23,24^. A remarkably slight difference (*p<0.01*) distinguished the marginally higher sensory S2 response latencies (*I_Per_* >0.25Bits, n=105, 33±15ms, ± S.D., Fig. 7B) from the expectedly fastest 3b neurons latencies (23 ± 9 ms, Fig. S6A). Analogously, S2 sensory coding latencies (*p<0.01*, 301 ± 71ms, Fig. 7B) were slightly slower than those from 3b neurons (241±45ms, Fig. S6B). Thus, sensory S2 neurons behave similarly to 3b neurons, with comparable latencies. Moreover, response (42 ± 21ms, Fig. S6A) and coding (359 ± 193ms, Fig. S6B) latencies for the whole population of S1 neurons (area 3b, area 1 and area 2), exhibit higher values than sensory S2 neurons (*p<0.01*). Neurons from DPC present even greater values: mean DPC latencies were 281 ± 190ms for response (Fig. S6A) and 484 ± 245ms for coding (Fig. S6B). Note that the whole population of S2 neurons displayed intermediate values between S1 and DPC. Shifting focus, categorical S2 neurons (*I_1s,P1_*>0.25Bits, n=150) show smaller, but comparable values of coding latencies (477 ± 201ms, Fig. 7C) than DPC neurons (*p<0.01*). Thus, DPC and categorical S2 neurons start coding P1 identity with relatively proportionate slow latencies. Even if the whole S2 neuronal population exhibit intermediate latencies^22^, sensory S2 neuron responses resemble those of 3b neurons and categorical S2 neuron coding resembles the slower trends found in DPC.

### Hierarchical intrinsic timescales across S1, S2 and DPC

In a recent work, timescales of intrinsic fluctuations in spiking activity across cortices were presented within a hierarchical framework^5^. This inherent timescale was measured using the autocorrelation function. We applied the same metric independently to the whole S2 population, as well as the identified sensory and categorical subgroups (Fig. 8A). We averaged the autocorrelation across the different groups of neurons, and we fit an exponential function to the average values (Methods). Surprisingly, we observed analogous autocorrelation functions, with similar decay, for the whole S2 population (τ = 178 ms), for the sensory (τ = 182 ms) or the categorical neurons (τ = 187 ms). Thus, even if the sensory and categorical neurons exhibit completely different response latencies, their autocorrelations show similar values (less than 3% difference, see Fig. 8A). Based on this finding, we speculate that even if each S2 subpopulation were engaged in antagonist roles, they are fundamentally embedded within the same cortical hierarchy.

**Figure 8.**
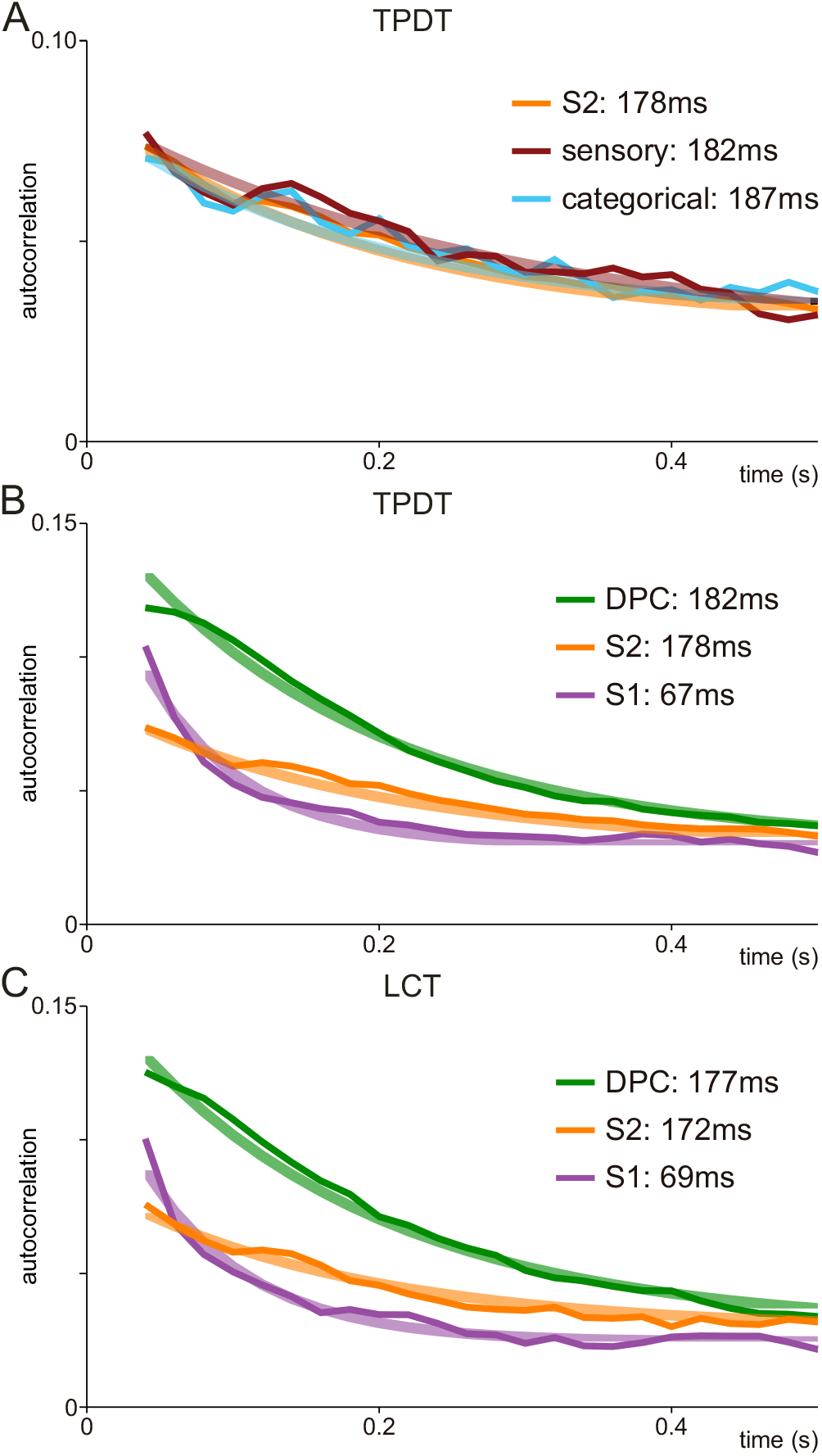
Spike count autocorrelation during basal periods. The autocorrelation function was computed for neuronal activity during the basal periods for the TPDT and LCT with 50 ms time bins. An exponential decay was fit to the autocorrelation function (Eq. 11). Thin light traces show the autocorrelation values averaged across each population of neurons. Wide dark traces display the exponential fit for each population. (A) Autocorrelation function for the entire S2 population (orange, n=1646, τ=178ms), S2 sensory population (brown, *I_Per_*>0.25 bits, n=105, τ=182ms) or S2 categorical population (blue, *I_1s,P1_*>0.25 bits, n=150, τ=187ms). Autocorrelation invariant to S2 subpopulations. (B) TPDT autocorrelation function for S2 whole population (orange, n=1646, τ=178ms) is compared against S1 population (purple, n=497, τ=67ms) and DPC population (green, n=1574, τ=182ms). Cortical areas exhibit a hierarchical order based on the autocorrelation function. (C) LCT autocorrelation function for S2 whole population (orange, n=313, τ=172ms) is compared against S1 population (purple, n=319, τ=69ms) and DPC population (green, n=462, τ=177ms). Autocorrelation functions analogous in both task conditions (TPDT and LCT).

To further substantiate these findings, when we implemented the same measure across S1 and DPC populations, our results established analogous hierarchical order as that proposed previously^5^ (Fig. 8B). As one would predict, autocorrelation functions from S1 neurons exhibit a much shorter decay constant, τ (τ = 67 ms). Hence, sensory information does not reverberate within the S1 network itself. Contrary to that, the DPC autocorrelation functions display greater values with a larger decay constant (τ = 182 ms). Even if the τ values are comparable among the S2 and DPC populations, the autocorrelation from DPC starts from much higher values. Notably, the autocorrelation functions across these different cortical areas were unaffected during the LCT (Fig. 8C). Even if the dynamic coding changed completely in S2 during the LCT (Fig. 3) and DPC coding completely disappeared (Fig. S7F), the autocorrelation functions did not change among conditions. These results are further evidence that autocorrelation is a measure of an inherent feature of each cortical network that does not depend on subpopulation or context. Even if there is a clear tendency, more research should be done in the future to clarify if the intra-area reverberation serves as an accurate component to establish the hierarchical order found here.

### Dynamical coding across areas S1, S2 and DPC during the TPDT and LCT

We reanalyzed recordings from S1 and DPC during the TPDT and the LCT^3^, and we computed the coding dynamics exactly as we did for S2 (Fig. S7). Notably, coding dynamics changed completely across cortices during the TPDT (Fig. S7A-C). While 3b (S1) activity is only involved in P1 or P2 coding identity during stimulation periods (Fig. S7A), S2 and DPC display much more complex dynamics. Whereas S2 neurons code P1 identity mainly during the early period of the working memory (Fig. S7B), DPC neurons exhibit strong persistent activity (Fig. S7C). These results are in conjunction with previous findings^22^. As we have described, S2 neurons exhibited high levels of P2 coding during the comparison. However, DPC neurons show robust class and decision coding. Note that P2 coding is negligible in DPC during the TPDT. Contrary to S2, DPC decision coding remains persistent during the last delay period (black trace). Therefore, during the comparison in the TPDT, S2 neurons code and categorize P2 in their activity, while DPC neurons were almost completely dedicated to comparison and decision coding.

As we have shown, S2 coding dynamics changed completely during the LCT. However, the coding responses in the DPC network were profoundly reduced to extinction (Fig. S7F). Analogous to sensory S2 neurons (Fig. 4F), 3b neurons remained invariant during this control task (Fig. S7D). Thus, information is coded in sensory S2 or 3b neurons independently of the task context. However, the duality of the dynamics in S2 vanished (Fig. S7E); only the sensory coding survived during LCT. A much more dramatic change was observed in the DPC population during the LCT (Fig. S7F); the rich and heterogenous coding dynamics observed in the TPDT ceased completely^3^. Therefore, task parameter information was inaccessible in this frontal area during LCT. As S2 population received the same sensory information but stopped categorizing during the LCT, we speculate that this area may play an important role in information flowing to frontal lobe areas. Our results suggest that S2 is presumably involved in this switch. To support this view, in a recent work in the visual system it was suggested that analogous parietal areas could be extremely relevant to ignition of higher order processes in cortical areas within the frontal lobe^25^.

## DISCUSSION

We examined the responses of S2 neurons and compared them with those responses from other cortices (S1 and DPC) during the TPDT and LCT. We empirically show conclusive evidence that demonstrates the coexistence of a duality in S2 response coding during the TPDT. On one hand, a substantial percentage of S2 neurons compose a sensory subgroup that is invariant to both behavior and cognitive demand of a tactile discrimination task. Alternatively, another specialized group of neurons categorically encode the stimulus identity with a clear dependence on task context (demanding or non-demanding) and behavioral reports (hits or errors). We elaborate on the significance and implications of these findings below.

Previous studies have proposed that S2 is connected to several cortical areas^26–32^. Hence, S2 appears to be located conveniently to receive both bottom-up and top-down inputs. Importantly, several works have suggested that S1 acts as a driver in the processing role of S2^11,12,33^, while others suggest that the somatosensory thalamus (VPL) plays the driver role instead^34^. Our results show strong evidence that a subgroup of the S2 population responds similarly to 3b (S1) neurons, with slightly longer average response latencies (33 ms versus 23 ms). However, whether these S2 responses are directly driven by 3b or VPL neurons is unclear. Admittedly, our experimental approach cannot address the origin of the sensory inputs received by S2, but this subgroup of neurons represented the stimulus patterns faithfully, in parallel to the faithful representation observed in area 3b of S1. Additionally, the S2 sensory responses are invariant to task context and behavioral responses, just as in area 3b of S1. Further, in other somatosensory tasks, we measured the response latencies of VPL neurons^35^, which demonstrated responses that arrived slightly earlier in VPL than in 3b. Another possible hypothesis is that S2 sensory neurons could depend on both VPL and S1 inputs. Nonetheless, for this uncertainty to be resolved, more experiments are required. However, it is important to mention that S1 is essential for performance of any somatosensory task, since when a lesion was applied to S1, the monkey’s ability to perform any tactile task was entirely ablated^36^. Whether or not S1 plays a driving role for S2, the role S2 plays in the processing network must depend in some manner on the sensory inputs from S1.

Although S2 sensory neurons could depend on the S1 inputs, much fewer neurons in S2 than S1 were entrained by the flutter stimulus. Further, a subgroup of S2 neurons exhibit complex and categorical responses that were not observed in 3b neurons^3^. Additionally, it is possible to find neurons with higher activity for the grouped or for the periodic pattern (dual coding)^37^. This dual and categorical representation emerges with similar latencies in S2 (~477 ms) and DPC neurons (~ 484 ms). These responses contrast sharply with the sensory S2 responses. Markedly, the categorical responses could represent the conversion of the sensory inputs, which are further distributed to areas downstream of S2. Changes in the representations from S1 to S2 have also been reported using a vibrotactile frequency discrimination task^2^ and employing texture surfaces^10^. Importantly, several neurons in S2 sustained their response coding during the early delay period between P1 and P2. This early working memory coding is markedly affected in error trials. Thus, while the sensory inputs are the same, the categorical transformation is directly correlated with the monkeys’ behavioral reports. Note that these units behave analogously to the early working memory neurons found in the DPC. These early DPC neurons also code P1 during the first few seconds delay period between P1 and P2. In contrast to DPC neurons, S2 neurons did not exhibit persistent coding during the working memory delay^2,22^.

We point out that the percentage of neurons that code class or stimulus pair combinations observed in the S2 population depends on information about P1, which must be accessible during P2. Thus, any comparison during P2 requires P1 information stored in memory (P2 = P1 or P2 ≠ P1). The percentage of neurons and variance that vanished at the beginning of the delay, with respect to P1, increased at the end of the delay and at the beginning of the P2 period. The disappearance of activity means that there was no persistent activity, which was confirmed by our analyses as well. The reappearance of activity associated to P1 requires that P1 information be fed-back to S2 neurons from a distinct area. We speculate that the information is retrieved from higher-order functioning cognitive areas, such as DPC, although there are several potential frontal lobe areas that could perform this feedback operation. These top-down signals between frontal areas may play a fundamental role in processing sensory information in S2.

It is important to emphasize that pure sensory and pure categorical neurons are the extreme cases of the S2 neural responses. Instead, the vast majority of S2 neurons exhibit intermediate dynamics (green points in Fig. 4C). One of the more intriguing results is that during the cognitively non-demanding control task, the LCT, the categorical responses either decreased or ceased, while the sensory information increased across the whole population; although not limited to, this included the intermediate cells. We speculate that the dual role of these neurons may be crucial in transforming pure sensory signals to more abstract responses. Again, further studies could illuminate to what extent abstraction occurs in S2, or if S2 receives a multitude of diverse signals and simply functions in a limited relay capacity. While either of these propositions may be possible, it is beyond the scope of our results to comment further.

We would like to highlight the high percentage of S2 neurons that continued coding the decision after the push button event. The emphasis we consider valuable from this type of response is that it acts during the period when the monkey has come to understand the result of his evaluative decision. Population variance associated with the categorical decision report also exhibited high values during this period. Moreover, this decision response is observed in both categorical and sensory neurons. To our knowledge, these are novel responses that have yet to be characterized in S2. This decision signal was lost during the LCT, which means that the signal could not be associated with a future motor plan; it is also a crucial result that the same signal and signal loss was observed previously in DPC neurons. Although the functional purpose of this categorical decision coding is not evident, we hypothesize that it could be associated with the reward signal observed afterwards in this S2 population. Both signals may be necessary for network re-wiring, employing the choice outcome to adapt future decisions. We are enthusiastic to see future studies that investigate this provocative potential role, both illuminating the dynamical function of S2 and identifying network operators that allow our monkeys to learn and perform these tasks.

Recent studies have presented evidence that the intrinsic timescale of neural fluctuations, estimated with the autocorrelation decay constant (*tau*, τ), increase from sensory to frontal lobe cortices^5^. Importantly, this result was found averaging across neurons from each cortical area during different cognitively demanding tasks. We found that this inherent timescale did not depend on the type of S2 response. Irrespective of their signals or coding function, the same autocorrelation functions occurred within the entire S2 population, which reveals one piece of insight: the subgroups are inherently parallel. This result suggests that this measure is indicative of a structural feature of each cortical area. Although timescales increase as a function of hierarchical stage (S1, S2 and DPC), they remain similar in the same cortical area during both the TPDT and LCT, even in the extreme case of losing all significant coding in the DPC neurons during the LCT.

Recurrent networks have been employed recently to model several responses observed in neural dynamics^38–40^. The coexistence of both sensory and categorical neurons in the same cortical area (S2) may represent an important constraint for new studies. In particular, the disappearance of the categorical responses during the control task and its relationship with behavior could provide challenging features to model. How does the network dynamic avoid categorizing the sensory inputs during the LCT? What are the roles of bottom-up signals in modelling these population dynamics? Our hopes are that future models will be able to inform the most confounding questions we have encountered through this study: how many roles does S2 play and how does it interact with a diverse array of inputs?

Recently, it was proposed that consciousness is related to an all-or-none distribution of neuronal responses across the brain^25,41–44^. Further evidence was given by the results obtained with a Global Neuronal Workspace (GNW) that simulates and compares the conscious state against a non-conscious state^41,45^. In this model, activity surpassing a threshold leads to an ignition where information is distributed across cortices. Importantly, if the subject does not attend to the stimulus, the ignition may fail. Applying this theory to our S2 findings, we speculate that this area could play a relevant role in broadcasting categorical information to frontal areas. Note that a portion of the responses observed in S2 neurons code task parameters in an abstract manner, and correlate with behavior (hit and error). The same effect was observed in DPC neurons^3^, with comparable latencies; however, during the LCT, these categorical responses in both S2 and DPC neurons ceased. Importantly, the sensory responses were not found in DPC under any task condition. Furthermore, unpublished results from our lab have suggested that animals with lesions applied to S2 are no longer able to perform the task adequately. Ultimately, this could mean S2 is a necessary component to the cognitive processing within the cortical network^46^.

To conclude, we would like to emphasize that both sensory and categorical responses were found within S2. While categorical responses ceased during the non-demanding task (LCT), the sensory responses remained unaltered. While categorical responses covaried with behavior, again the sensory were invariant. A subgroup of S2 neurons behaved like area 3b neurons (S1), while another subgroup mirrors the behavior found in DPC neurons, forfeiting their coding response during the non-demanding task. We speculate that this area may play a fundamental role in transforming sensory inputs to more abstract, conceptual and categorical responses; the role of S2 could even be that of a gateway to downstream frontal areas^1^. Therefore, S2 may act as a switch network: it is always receiving the same sensory inputs, but selectively converting and transmitting abstract representations when the task demands it. This may be a central processing principle, not only for S2, but also for other parietal areas related to other sensory modality tasks.

## ACKNOWLEDGEMENTS

We thank H. Diaz and G. Diaz de Leon for technical assistance. The research of R.R. was partially supported by the Dirección General de Asuntos del Personal Académico de la Universidad Nacional Autónoma de México (PAPIIT-IN210819).

## MATERIALS AND METHODS

### Temporal pattern discrimination task (TPDT)

The TPDT used here has been previously described^3^. In brief, two monkeys (*Macaca mulatta*) were trained to report whether the temporal structure of two vibrotactile stimuli patterns (P1 and P2) of equal mean frequency (5 Hz, 5 pulses) were the same (P2 = P1) or different (P2 ≠ P1; Fig. 1A). The temporal structure of each pattern was either grouped (G) or extended (E) with a fixed stimulation period of 1s. The 5 pulses were delivered periodically during extended pattern (E) or 3 grouped centered pulses with a smaller distance between during grouped pattern (G). Monkeys performed the task in blocks of trials in which the two stimulus patterns had a fixed mean frequency. The right arm, hand and fingers were held comfortably but firmly throughout the experiments. The left hand operated an immovable key (elbow at ~90°) and two push buttons in front of the animal, 25 cm away from the shoulder, at eye level. Stimuli were delivered to the skin of one digit from the distal segment of the right, restrained hand via a computer-controlled stimulator (2 mm round tip, BME Systems, Baltimore, MD). The initial event marks the beginning of the trial by descending the probe to a skin indentation of 500μm (probe down, “pd” in Fig. 1A). Vibrotactile stimuli consisted of trains of short mechanical pulses; each pulse consisted of a single-cycle sinusoid lasting 20ms. Time is always referenced to first stimulus onset (0s corresponds to the start of P1). In a trial, P1 and P2 were delivered consecutively to the glabrous skin of one fingertip, separated by a fixed inter-stimulus delay period of 2s (1 to 3s). Each stimulus could be one of the two possible patterns: grouped (G, upper trace of Fig. 1A) or extended (E, lower trace of Fig. 1A) pulses. Therefore, in total there were four possible P1-P2 combinations, denominated classes: G-G (class 1, c1), G-E (class 2, c2), E-G (class 3, c3) and E-E (class 4, c4). These were presented in pseudo-random order to the monkeys across trials. The monkeys were asked to report whether P2 = P1 (match: combinations E-E and G-G) or P2 ≠ P1 (non-match: combinations E-G and G-E) after a fixed delay period of 2s (4 to 6s) between the end of P2 and the mechanical probe rising from the skin (probe up event, “pu” in Fig. 1A). The “pu” was the go signal that triggered the animal’s release of the key (“ku” in Fig. 1A). The monkey indicated their decision by pressing one of two push buttons with the left hand (“pb” in Fig. 1A, lateral push button for P2 = P1, medial push button for P2≠P1). Because the two stimulus patterns had equal mean frequency over their full duration (1s), the decision had to be based on comparison of their temporal structure. The animal was rewarded for correct decisions with a drop of liquid. Animals were handled in accordance with standards of the National Institutes of Health and Society for Neuroscience. All protocols were approved by the Institutional Animal Care and Use Committee of the Instituto de Fisiología Celular, Universidad Nacional Autónoma de México.

### Light control task (LCT)

During this task, events proceeded exactly as described above and in Fig. 1A, except that when the probe touched the skin (“pd”), one of the two push buttons was illuminated, indicating the correct choice. The mean stimulus frequency was held constant as well (5Hz). The monkey grasped the key until the probe was lifted, but in this case the light was turned off when the probe lifted from the skin. The monkey was rewarded for pressing the illuminated button. Maintaining stimuli and arm movements identical to the TPDT, the decision must be based on the visual stimuli instead.

### Task design and performance

The TPDT is not a simple variation of the vibrotactile frequency discrimination task (VFDT,^47^). Some cognitive demands and the basic structure of the tasks are similar: both require attention to two separate vibrotactile stimuli (TPDT: P1, P2; VFDT: f1, f2), working memory and a comparison to reach the decision report. Nevertheless, the TPDT requires a very different transformation of the stimuli; as they only differ by their temporal structure, any computation must be restricted to the internal structure to identify, categorize and distinguish between them^3^. Further, the comparison process is significantly different between the two tasks. Expanding on the necessitated computation, the VFDT can be solved by computing a difference between the parametric representation of the stimulus frequencies to indicate whether f1 > f2 or f1 <f2, whereas the TPDT offers no comparable method of solution (in any trial P1 and P2 always have the same mean frequency). The TPDT demands a match (P2 = P1) vs. non-match decision (P2 ≠ P1). Hence, the comparison employs categorical representations (instead of parametric) of the stimulus patterns.

We computed the average performance across S2 recording sessions (84±7%, Fig. 1B). Although each animal received between two and three years of training, this task was difficult enough not to allow 100% performance; this reflects the very high cognitive demands of the TPDT. To provide some context, the average training period to achieve similar performance levels for the VFDT was about six to eight months^48^; for the vibrotactile detection task^49^, the average time was two months. After training in the TPDT, the monkeys saturated their average performance around 84% (Fig. 1B, n=423 recording sessions). In addition, the performance was statistically identical for each class^3^. Notably, task repetition across recording sessions did not improve performance. However, the performance for the LCT was consistently 100% (Fig. 1B, n=76 recording sessions); this reflects the lack of cognitive demand required for the guided-task, as intended by design. As a final observation, the animals were first trained in the LCT, and then gradually introduced to the TPDT. During the recording sessions in S2 (Fig. 1C), animals switched between performing the TPDT and the LCT.

### Recordings

Neuronal recordings were obtained with an array of seven independent, movable microelectrodes (2-3 MΩ^4^) inserted into S2 (Fig. 1C), either contralateral (left hemisphere) or ipsilateral (right hemisphere) to the stimulated hand. We collected neuronal data in blocks using different mean frequencies^3^. However, for the analysis described below we will focus on the neuronal responses with the stimulus set illustrated in Fig. 1A (5 Hz). In general, we recorded 20 trials per stimulus pair (c1; c2; c3; c4). Recording sites changed from session to session; the locations of the penetrations were used to construct surface maps in S1, S2 and DPC by marking the edges of the small chamber (7 mm in diameter) placed above each area. In area 3b (S1) we recorded neurons with cutaneous receptive fields confined to the distal segments of the glabrous skin of one fingertip of digits two, three or four. All recordings in DPC were made in the arm region F2. This region is in front of M1 (F1), lateral to the central dimple, posterior to F7 and the genu of the arcuate sulcus^50^. Neuronal recording protocol was identical for both the TPDT and LCT.

### Datasets

We recorded 1646 S2 neurons using the TPDT stimulus set with 5 Hz mean frequency. Additionally, we have a dataset of n = 313 neurons that were tested in both the LCT and in TPDT using the 5Hz mean frequency set. These neurons were used to compare periodicity and firing rate information between the cognitively demanding TPDT and the guided LCT (Fig. S4).

For each of neuron of the datasets (n = 1646 and n = 313), we calculated a time-dependent firing rate per trial using a 200 ms deterministic square kernel with 50 ms steps, beginning 1s before stimulus pattern P1 until the end of the trial (1.5s after the push button press). Importantly, each dataset is defined by four dimensions: N, number of neurons; C, stimulus conditions (classes, always 4); T, time (−1 to 7.5s, always 170 bins); K, number of hit trials (for each class). Further, we constructed a similar dataset with error trials for the 5 Hz TPDT stimulus set. Each recorded neuron had on average 2.3 error trials for a given class. A remarkable feature of this task design is the low number of stimulus conditions (four classes), which were equally demanding for the subject. This design allowed us to have, on average, 17.7 hit trials (and 2.3 error trials) per stimulus class for each studied neuron.

### Single neuron coding

This analysis was designed to quantify whether the activity of single S2 neurons was modulated as a function of time by the four stimulus conditions (which we call classes) used in the task: c1 (G-G); c2 (E-G); c3 (E-G) and c4 (E-E). We employed the same coding scheme used previously to identify single neuron coding in DPC and S1^3^.

Employing only hit trials, we constructed a neuron firing rate distribution for each class. At each time bin we used the receiver operating characteristic (ROC) to identify class-differential responses; using these class firing rate distributions, we computed the area under the ROC curve (AUROC value) for the six possible class comparisons: c1 vs. c2; c1 vs. c3; c1 vs. c4; c2 vs. c3; c2 vs. c4; and c3 vs. c4. To determine significant AUROC values, we performed a permutation test by randomly shuffling the class labels across trials, while re-computing the AUROC values with the shuffled trials. If the unshuffled AUROC value (≠0.5) reached or exceeded the 95% of the distribution obtained from 1000 shuffled surrogates, responses for the two compared classes were labeled statistically different (p<0.05); otherwise, they were labeled as equal. We should emphasize that statistical equality means that there is not enough neuronal response information to differentiate the two distributions; this does not mean that both distributions were the same.

From this, we produced a library of binary words; for each 200 ms bin we had six digits resulting from the six comparisons. In this coding scheme, the 0’s are as important as the 1’s. The criterion to assign both was very strict: to avoid random assignments at each time window, we only assigned a binary label of statistical equality (0) or inequality (1) if the same digit was kept for at least four consecutive bins, otherwise no label would be assigned, and that time bin was excluded from the classification. This part of the coding scheme was designed to correct for multiple comparisons. It is important to note that for each time bin this procedure generates a unique code for each neural response, one of our “binary words”. However, we isolated four relevant response or coding labels from the 64 binary words. These four profiles are explained below. From 64 binary words, we isolated 7 associated to our labels, while the rest represent mixed or ambiguous codes. Using the binary words computed from the six AUROC values as described above, each time bin was tested for classification into one of four possible coding profiles during the TPDT and LCT.

#### P1 coding

This profile applied to responses that tracked the identity of the P1 pattern. In this case, the responses must be similar for classes c1 (G-G) and c2 (G-E), and for c3 (EG) and c4 (E-E), which have the same P1, but must differentiate between all other class comparisons, which have different P1 patterns.

#### P2 coding

As described above, but for responses that tracked the identity of P2. Responses must be similar for c1 and c3, and for c2 and c4, which have identical P2, and must be different for all other class combinations, which have different P2 patterns.

#### Class-selective coding

This profile corresponds to neurons that responded preferentially to one of the four classes. Time bins were labeled according to the class that selectively evoked a response. We associated four binary words with this profile, pursuant to a single rule: the preferred class evoked a unique response, while the three non-preferred classes were indistinguishable between each other.

#### Decision coding

In this profile, responses must be similar for classes c1 (G-G) and c4 (E-E), as well as c2 (G-E) and c3 (E-G), which share the same outcomes (either P1=P2 or P1≠P2) and differ for all other class comparisons with distinct outcomes.

Time bins where the six comparisons did not fit any of the binary words described above were considered to be non-coding. Further, to consider that a neuron had significant coding, a minimum 4 consecutive bins must maintain the same profile. Applying this procedure across all neurons allowed classification of encoding dynamics as functions of time (Fig. 3A and B, Fig. 4E and F). This coding scheme rendered two advantages: 1) being able to quantitatively assess and describe all the possible neural codes during all task epochs, and 2) generating coding types that would not overlap in their meaning.

### Instantaneous coding variances across the population

For each neuron, we averaged the time-dependent firing rate of hit trials per class (c1, c2, c3 or c4). Using the peri-stimulus time histogram (PSTH) of each neuron, we constructed pseudo-simultaneous population responses by combining neural data mostly recorded separately. For each time and class, the population response is defined by an N-dimensional vector in which each component represents the firing rate from a different neuron. This means that including all the recorded neurons (n = 1646), we obtained a 1646-dimensional firing rate vector that depended on the time and class 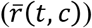). The population firing rate averaged over all hit trials 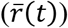) was an N-dimensional vector that measures the mean response for each neuron (*r^i^*(*t*)) as a function of time. For the LCT control condition, the population response was a 313-dimensional firing rate vector.

At each time point, the population instantaneous coding variance (*Var_COD_*, Figs. 3C and S3C-F, blue trace) was computed as the quadratic square sum of the firing rate fluctuations among classes and neurons:

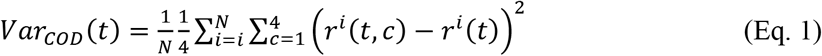

This metric, normalized per neuron, measures the population’s variation of firing rate between classes at each time point. In this case, *Var_COD_* will be associated to any class-related change in the population activity and to stochastic fluctuations (residual noise).

To evaluate the influence of each kind of coding on *Var_COD_*, we calculate the instantaneous variance associated to each task parameter. At each time bin, the population instantaneous P1 variance (*Var_P1_*, Figs. 3C and S3C-F, cyan trace) was computed as the quadratic square sum of the firing rate fluctuations among P1 identity and neurons:

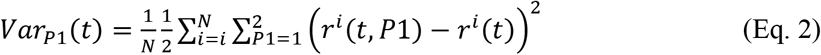

Analogously, the population instantaneous P2 variance (*Var_P2_*, Figs. 3C and S3C-F, light green trace) measures the firing rate fluctuations among P2 identity and neurons:

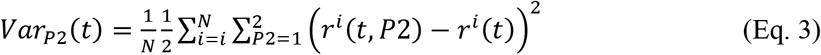

The population instantaneous decision variance (*Var_DEC_*, Figs. 3C and S3C-F, black trace) measures the firing rate fluctuations of decision identity and neurons:

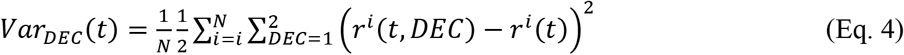

The value of *Var_Cod_* during the period immediately before P1 onset represented the inherent stochastic fluctuation (residual noise) in the firing rate estimates (~2[sp/s]2); to be interpreted as a degree of population coding, *Var_Cod_* should be higher than this resting state variance (basal variance). The same reasoning applies to the other specific variances. Accordingly, *Var_Cod_* and *Var_P1_* depart from their basal values at the same time bins (Fig. 3C). Further, the times at which any of the specific variances depart from their basal value coincide with the emergence of significant coding in individual neurons (compare Fig 3A with 3C and Fig. 3B with 3D).

#### Sensory population response

In order to describe the sensory population responses of S2 (n=105), we normalized the firing rates for each time bin (50 ms window displaced every 10 ms) using the z-score transform. The z-score was computed by subtracting from each trial (hit, error, and control trials) the mean firing rate and dividing the result by the standard deviation (SD) at each time window. The mean and SD for each neuron were calculated using the recorded firing rate activity in hit trials from all time bins in the interval from −1 to 7.5s of the task. We calculated a mean z-score value for hit, error and control (light control task, n=41) trials for each class to obtain an average sensory population response as a function of time. Finally, we transformed back the mean population z-scores to show responses in terms of firing rates instead of z-scores (Fig. 5). Back transformation was computed using the average firing rate values and SDs from all sensory neurons.

#### Firing rate information

Using the firing rate values, we measured their association with P1 and P2 in terms of Shannon’s mutual information:

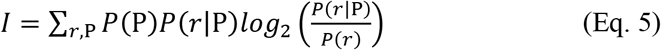

Here, the information (I), measured in bits, quantifies the accuracy with which the neural response (the firing rate *r*) can be used to determine the identity of the stimulus pattern (P). The expression *P*(*r*) corresponds to the probability of observing a response (*r*) regardless of the stimulus pattern; it was computed using the firing rate probability distribution from all hit trials during the same time window. *P*(P) represents the probability that the stimulus pattern takes a value P (G or E), considering only hit trials. *P*(*r*|P) is the conditional probability of observing a response *r* given a specific stimulus pattern P.

Importantly, to calculate the categorical information metric, we employed 1000 ms windows that cover the whole first stimulus (from 0 to 1s) or the whole second stimulus (from 3 to 4s). Then, we quantified the decodable firing rate information conveyed by each neuron about pattern identity (G or E) during P1 or P2, employing a 1s integration window (*I*_1*s*_):

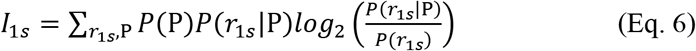

Note that *I*_1*s*_ in neurons that are tightly phase locked to stimulation pulses (phase-locked or sensory neurons), should be near zero (see Fig. 3 in^3^). Since the number of pulses is the same for each type of pattern G and E, if each pulse is represented equally by a sensory neuron, the firing rate during the whole stimulus period (1s) should be approximately the same. This means that 1s-firing rate mutual information (*I*_1*s*_) associated with the pattern identity is near zero for sensory neurons. Contrary to that, categorical neurons should have higher values of *I*_1*s*_, where pulses generated different responses depending on the pattern identity (G or E).

In Fig. 6, we computed the firing rate mutual information associated with the identity of P1 during hit or error trials for different subpopulations of S2 neurons. We z-scored the 200 ms firing rate responses from each hit or error trial at each time bin. Then, we joined the z-score values from different neurons to calculate the population z-score conditional probabilities (*P*(*z*(*t*)|P1)) associated with each pattern (E or G). Note that we constructed different distributions for hit and error trials. Next, we used the z-score population probabilities to estimate, per time bin, the mutual information associated with P1 during hit and error trials:

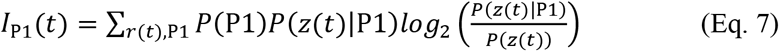

Analogously, we calculated the population firing rate mutual information associated with the decision identity during hit or error trial (Fig. S5B). As before, we computed the z-score normalization to the 200 ms firing rate responses, splitting hit and error trials. Then, we constructed population probability distributions associated to decision identity (equal or different) during hit or error trials:

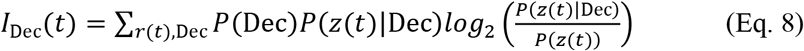

Finally, we estimated the firing rate mutual information associated with reward (Fig. S5C). As explained above, at each time bin, we computed population probability distributions, condensing the 200 ms z-score activity from different trials and neurons. In this case, we computed a distribution with all hit trials and another with all error trials. We employed these two population probability distributions to calculate the amount of information associated with the rewards, conveyed in the firing rate of the population:

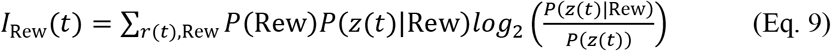

#### Periodicity information

The extended patterns (E) are periodic with a temporal spacing of 240 ms between pulses (freq=4.16Hz). Contrary to that, grouped patterns (G) are aperiodic; there are three centered grouped pulses (Fig. 1A, up pattern). Based on the temporal stimulus structure, a phase-locked neuron (sensory) should respond periodically during an extended pattern (E) at 4.16 Hz but not during grouped patterns (G). We aimed to compute the mutual information associated to the pattern identity (G or E) that is conveyed by the periodicity of the neural responses. To accomplish that, we employed Fourier decomposition of the time signals formed by the evoked trains of spikes during stimulation periods. For each trial, the power spectrum of the spike train evoked during stimulation (first pattern P1, or second pattern P2) was computed and normalized. We removed the DC component, so that the total power summed over all positive frequency bins was 100%^2,51^. Employing this methodological approach, the number of spikes contained in each train had little effect on the resulting Fourier amplitudes, which indicate the proportion of power for each frequency bin. Then, Fourier amplitudes were mainly determined by the temporal arrangement of the spikes, not by their number. Each trial was first transformed to firing rate employing a quadratic and deterministic kernel of 24ms and 0.6 ms step. The width of the frequency bins was 0.97Hz. This value was limited by the duration of the stimulation period, which for the Fourier analysis we took as 1228.8ms. This means that for each stimulus period we employed 2048 points to compute the Fourier transform, starting 50ms before and finishing 178.8ms after P1 or P2 periods.

From each trial, we extracted the two power spectra values associated to the two Fourier frequencies (3.88Hz and 4.85Hz) that are nearest to the periodic stimulation frequency (freq=4.16Hz). These values should increase for evoked spikes that are more tightly phase-locked to the periodic stimulation pulses. Suppose a neuron is strongly phase-locked to the periodic pattern (E) and fires spikes somewhat like a clock, one or two spikes per stimulation pulse, in an approximately periodic fashion. In its spectra, the maximum power would be at the periodic pattern frequency (freq=4.16Hz). Hence, for a sensory neuron, these values should be high during E patterns and small during G patterns.

As we explained for the firing rate, the mutual information that the periodicity of the response at 4.16Hz (*pf*) provides about the stimulus pattern (P) is calculated from the probability distributions relating these two variables. The function *P*(*pf*|P) represents the conditional probability of observing a spectrum value at 4.16Hz given that the stimulus pattern had a value of P (G or E). The expression *P*(*pf*) describes the probability of observing a spectrum value at 4.16Hz regardless of the value of the stimulus pattern, and *P*(P) is the probability that the stimulus takes a value of P (G or E). Then, the information that the spectrum value at 4.16Hz provides about the pattern identity can be computed as:

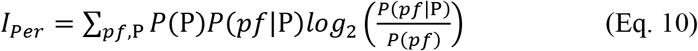

For all the mutual information values computed across this work (Eqs. 5–10), a correction for sampling bias was applied^52^. Furthermore, the significance of mutual information values for neurons labelled as sensory and categorical was computed through a permutation test, with the significance criterion set to the *p<0.01* level.

#### Choice probability

The choice probability index (CPI) was calculated using methods from signal detection theory^53^. In this case, the ROC measures the overlap between hit and error responses for each stimulus pair (P1, P2). A value of 0.5 indicates full overlap, whereas 1 and 0 indicate no overlap between distributions. Thus, the CPI quantifies the selectivity for one or the other decision outcome during the discrimination process. To compute the CPI as function of time, we used a window of 200 ms duration moving in steps of 50 ms, beginning at P2 and ending 1500 ms after the animal reported the comparison between P2 and P1. To combine the responses from all neurons at each time bin, the CPI values were averaged across all S2 neurons with decision coding (Fig. S5A).

#### Response latencies

We calculated two different latencies, a response latency, which corresponds to the time at which the stimulus-driven neural activity (during P1) becomes significant, and a coding latency, which corresponds to the time at which the encoded signal becomes significant (during P1).

##### Response latency

Firing rate distributions were generated at each time point using a time window of 200 ms sliding steps of 1 ms during P1, and were compared against the rates obtained in a control period (200 ms before P1 onset) using the ROC method^22,53^. The first time bin at which the AUROC was significantly different from 0.5 (permutation test, p < 0.05) for five consecutive bins was considered as the response latency to P1.

##### Coding latency

This latency varied depending on the coding profile of the cells. The P1 coding latency was estimated for each neuron by identifying the first of three consecutive bins significantly coding patterns G or E.

#### Autocorrelation analysis

The autocorrelation functions of spike counts were computed following the same methodological procedure as in^5^ for single neurons. The basal period (−1 to 0 sec) was divided into non overlapping, successive time bins of 50ms duration. Then, for two time bins separated by a time lag *t*, we calculated the across-trial correlation between spike counts *N*. Next, we averaged the correlation values computed for each neuron and time lag *t* across the population. Afterwards, this averaged population autocorrelation function of the time lag *t* between bins was fit by an exponential decay with an offset:

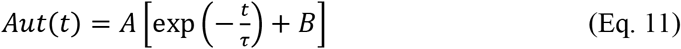

In this equation the autocorrelation tau (τ) measures an intrinsic population timescale. The offset (*B*) represents the contribution of timescales much longer than our observation window. We fit Eq. 10 to the full autocorrelation data from all neurons and trials. Hence, fits were performed at the population level rather than single-neuron level. To be able to fit this equation to the single neuron level, much more recorded trials per cell are required. To fit Eq. 10 to the population autocorrelation data, a nonlinear least-squares fitting via the Levenberg-Marquardt algorithm was employed.

---

**Figure S1.**
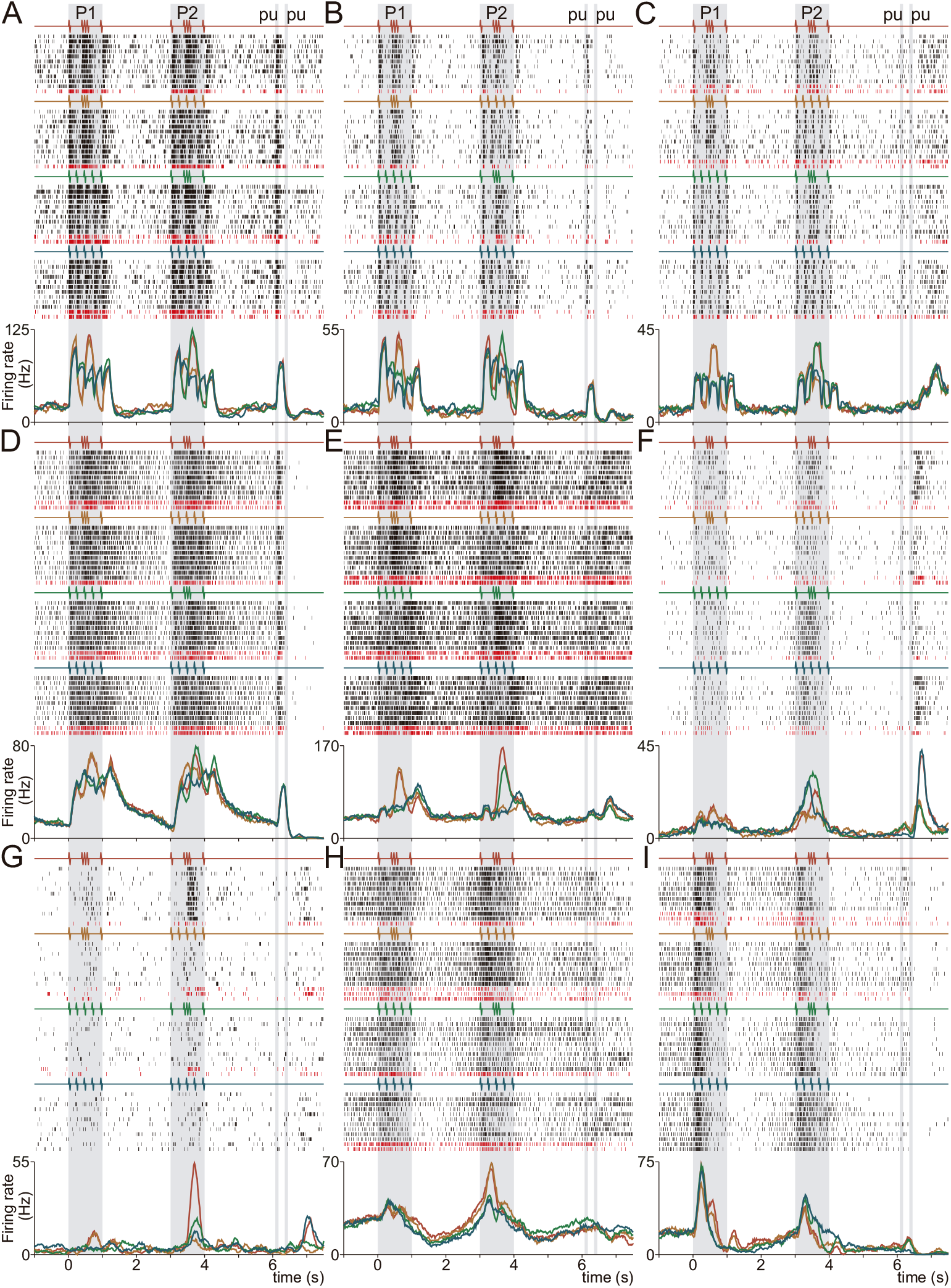
Single neuron activity in S2. S2 neurons display sensory (A-B), intermediate (C-E) and categorical (F-I) responses during TPDT. (A-I) Raster plots of nine exemplary S2 neurons sorted according to the four pattern pairs of G and E stimuli, delivered during P1 and P2. Each row is one trial, and each tick is an action potential. Trials interleaved randomly, blocks sorted by classes (only 10 out of 20 trials per class are shown). Correct and incorrect trials are indicated by back and dark red ticks, respectively. Traces below the raster plots are firing rate averages per-class [peristimulus time histograms (PSTHs)] for each neuron. Each color refers to one of the four possible pattern pairs of G and E: c1 (G-G, red), c2 (G-E, orange), c3 (E-G, green) and c4 (E-E, blue).

**Figure S2.**
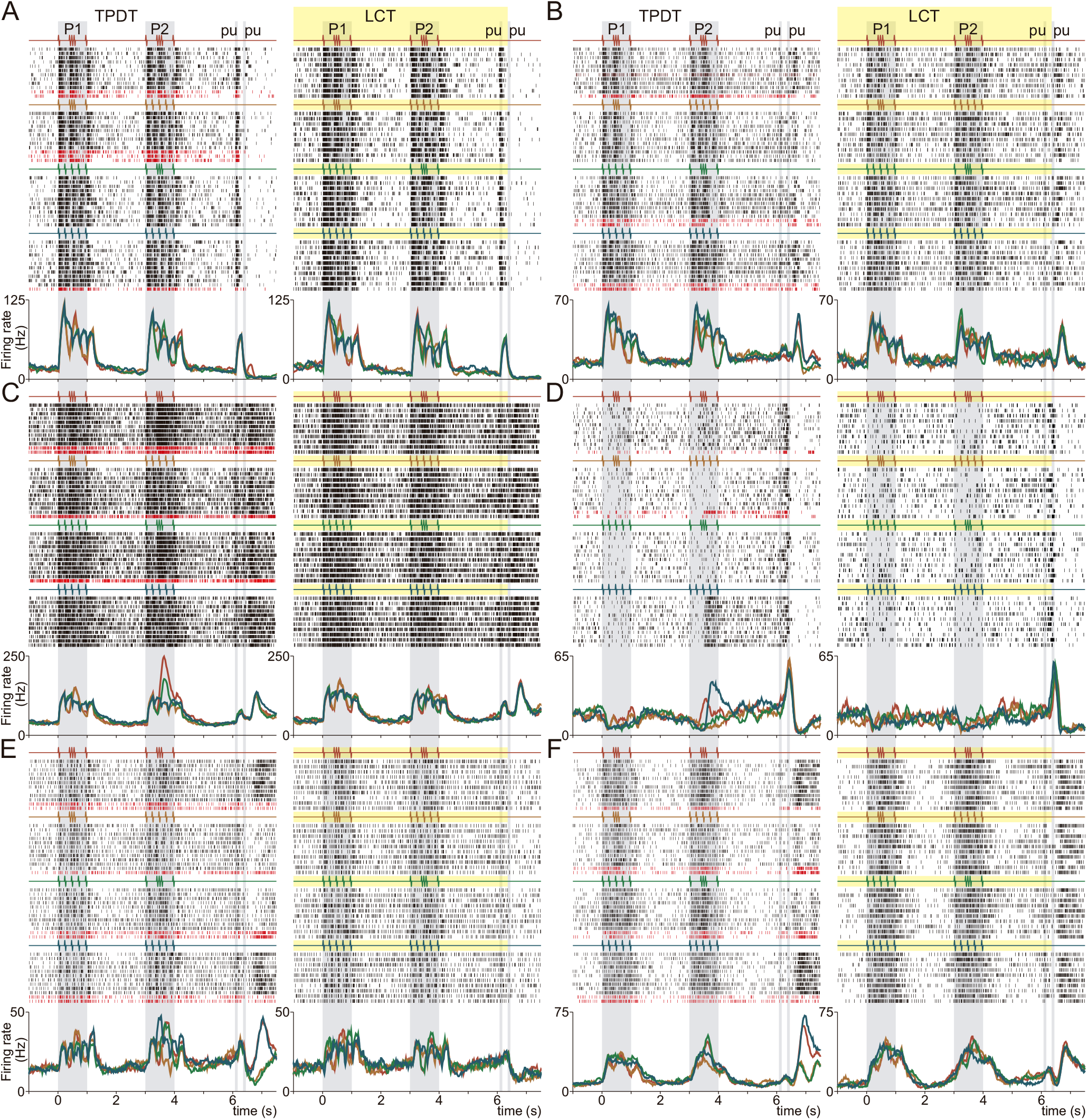
Single S2 neurons activity during the TPDT and LCT. S2 sensory responses remained unaltered during TPDT and LCT. S2 categorical responses diminished during LCT. (A-F). Raster plots of six additional exemplary S2 neurons tested in both experimental conditions: TPDT (left panels) and LCT (right panels). Responses are sorted according to the four possible combinations of G and E stimulus pattern delivered during P1 and P2. Correct and incorrect (only in TPDT) trials are indicated by black and dark red ticks, respectively. Traces below the raster plots are average per-class firing rates (PSTHs) for each neuron. Each color refers to one of the four possible stimulus combinations of G and E; the resulting four classes are: c1 (G-G, red), c2 (G-E, orange), c3 (E-G, green) and c4 (E-E, blue).

**Figure S3.**
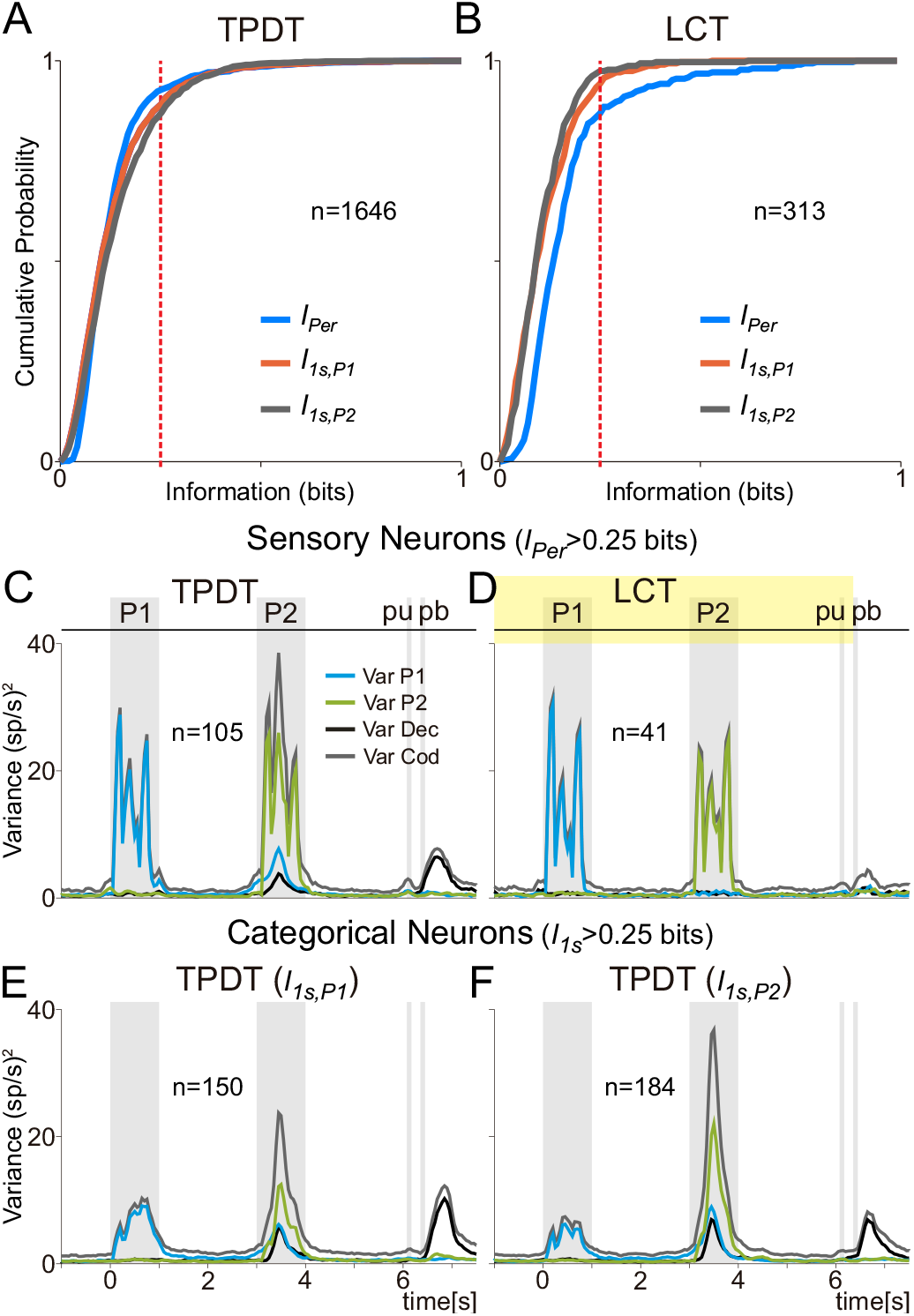
Mutual information and population variance in sensory and categorical neurons. (A-B) Cumulative probability distribution for periodicity (*I_Per_*, blue trace, Eq. 10), 1s firing rate mutual information about P1 (*I_1sP1_*, orange, Eq. 6) or P2 identity (*I_1sP2_*, grey, Eq. 6). The value on the y axis represents the fraction of neurons with mutual information smaller or equal to the amount indicated on the x axis. (A) Population of neurons recorded during the TPDT (n=1646). (B) Neurons recorded during LCT (n=313). Note that a greater percentage of neurons exhibit higher *I_Per_* during LCT than TPDT. The red dashed lines indicate the mutual information limit (*I*>0.25bits) used to label S2 neurons as sensory or categorical. This arbitrary boundary is useful to explore the dynamics features of the S2 neurons with higher amounts of *I_Per_* or *I_1sP1_*. (C-F) Population instantaneous coding variance (*Var_COD_*, grey, Eq. 1), P1 variance (*Var_P1_*, cyan, Eq. 2), P2 variance (*Var_P2_*, green, Eq. 3), and decision variance (*Var_Dec_*, black, Eq. 4) computed in different subpopulation of S2 neurons. (C) Sensory neurons (*I_Per_*>0.25 bits, n=105) during TPDT. Most coding variance is related to stimulus phase-locked activity. (D) Sensory neurons (*I_Per_*>0.25 bits, n=41) during LCT. (E-F) Categorical neurons computed during P1 (left, *I_1s,P1_*>0.25 bits, n=150) or P2 (right, *I_1s,P2_*>0.25 bits, n=184). Note the marked differences between variance computed with sensory and categorical neurons during the TPDT.

**Figure S4.**
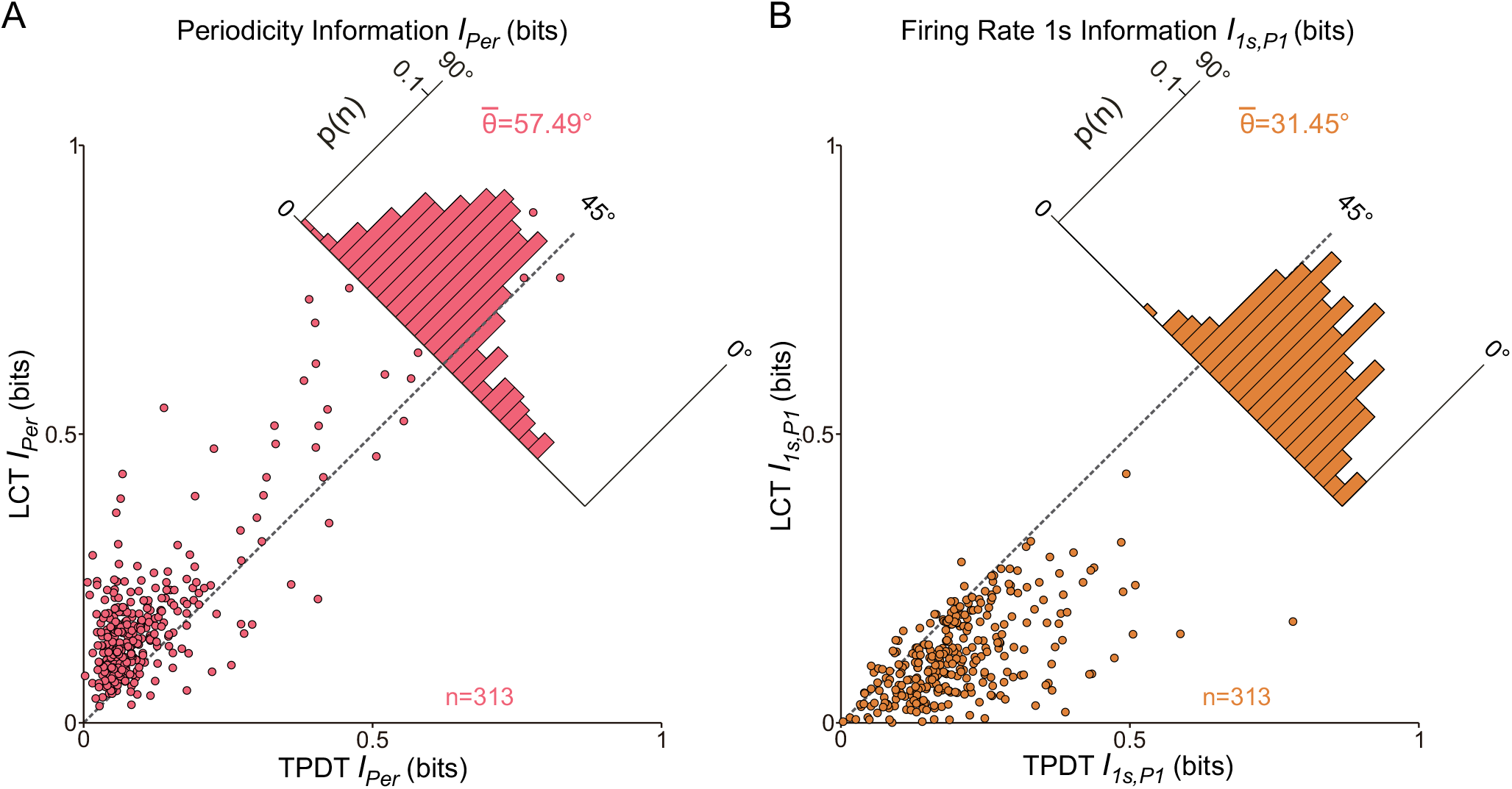
Single neuron periodicity and categorical information between TPDT and LCT. In this figure, we restricted our analysis to the neurons that were recorded during both TPDT and LCT (n=313). Each dot in these plots corresponds to one neuron tested in the two different task conditions. (A) *I_Per_* in TPDT (x axis, Eq. 10) is compared with *I_Per_* in LCT (y axis). Inset histogram show angular distributions for the population (<θ>=57.49°). On average, *I_Per_* tend to be higher during LCT than TPDT. (B) *I_1sP1_* in TPDT (x axis, Eq. 6) is compared with *I_1sP1_* in LCT (y axis). Inset histogram show angular distributions for the population (<θ>=31.45°). For the same group of neurons, *I_1sP1_* tend to be higher during TPDT than LCT.

**Figure S5.**
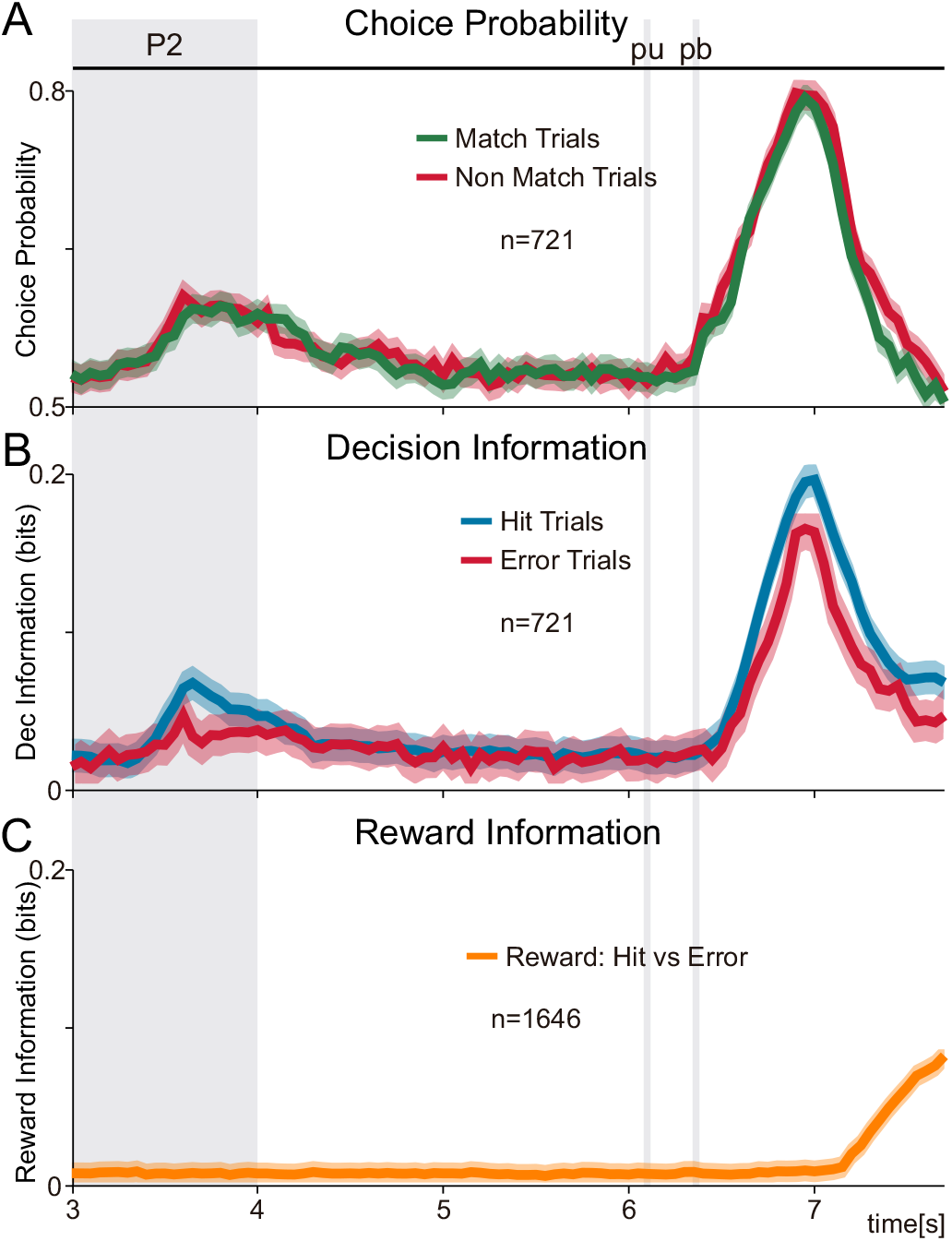
Choice probability, decision and reward mutual information during hit and error trials. (A) Mean choice probability (CP) as function of time for neurons (n=721) with at least 3 consecutive time bins with decision coding. Higher CP values were obtained after push button (pb). (B) Population firing rate mutual information associated with decision outcome computed as function of time (*I_Dec_(t)*, Eq. 8) during hit (blue) and error (red) TPDT trials for the same S2 neurons than in the previous panel (n=721). Hit and error trials give comparable amount of decision information after pb. (C) Population firing rate mutual information associated with reward computed as function of time (*I_Rew_(t)*, Eq. 9). Population probability distributions with all hit or error trials were employed to estimate the amount of mutual information associated with reward, conveyed in the firing rate of the population. *I_Rew_(t)* increased after animals were rewarded for correct decision (<*t_Rew_*>=6.92s). Shadows indicate the information or CP confidence intervals at 95%.

**Figure S6.**
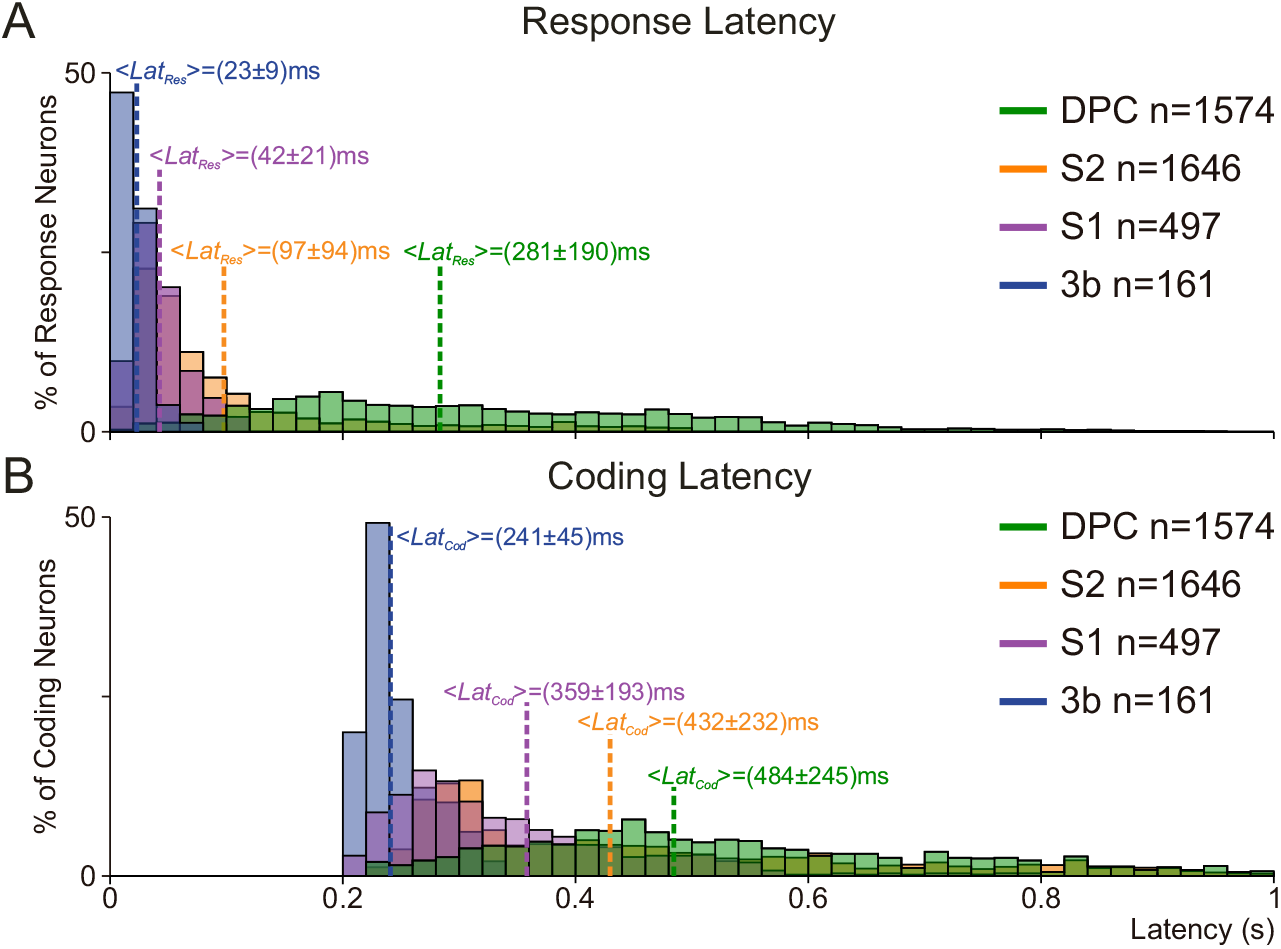
Response and coding latencies across cortical areas. (A-B) Response (*Lat_Res_*, top) and coding (*Lat_Cod_*, bottom) latency distributions are computed during TPDT for the whole dorsal premotor cortex (DPC) population (green, n=1574), the whole S2 population (orange, n=1646), the whole somatosensory cortex (S1) population (purple, n=497) and the whole area 3b (S1) population (blue, n=161). Mean values for each latency distribution are indicated with dashed lines (± values indicate standard deviation, S.D.). Note that neurons from area 3b are also included in S1 population. A cortical area hierarchical order could be established in the response and coding latencies.

**Figure S7.**
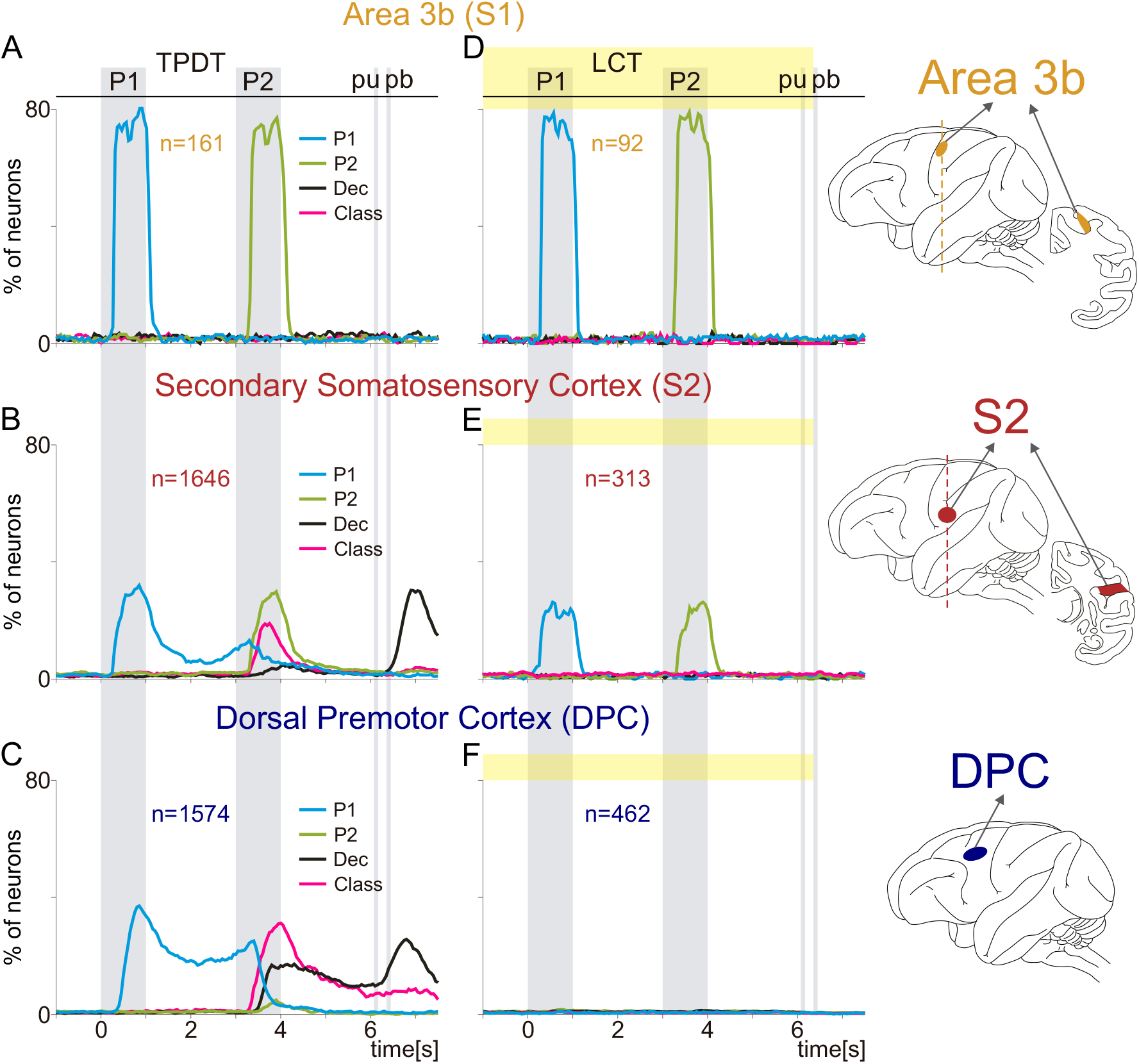
Population coding dynamics across cortex during TPDT versus LCT. Percentage of neurons with significant coding as a function of time during TPDT (right, n=1646, A-C) or LCT (left, n=313, D-F). Traces refer to P1 (cyan), P2 (green), class (pink, stimulus pair combinations), and decision coding (black). Neurons were recorded from different cortical areas: Area 3b (TPDT [A, n=161] and LCT [D, n=92]); S2 (TPDT [B, n=1646] and LCT [E, 313]); DPC (TPDT [C, n=1574] and LCT [F, n=462]). Note that Area 3b is included in the first somatosensory area (S1). The same y scale (0 to 80%) was used across panels to facilitate comparison (same scale used for Fig. 4E-H).

